# Connectome Spatial Smoothing (CSS): concepts, methods, and evaluation

**DOI:** 10.1101/2021.09.19.461011

**Authors:** Sina Mansour L., Caio Seguin, Robert E. Smith, Andrew Zalesky

## Abstract

Structural connectomes are increasingly mapped at high spatial resolutions comprising many hundreds—if not thousands—of network nodes. However, high-resolution connectomes are particularly susceptible to image registration misalignment, tractography artifacts, and noise, all of which can lead to reductions in connectome accuracy and test-retest reliability. We investigate a network analogue of image smoothing to address these key challenges. Connectome Spatial Smoothing (CSS) involves jointly applying a carefully chosen smoothing kernel to the two endpoints of each tractography streamline, yielding a spatially smoothed connectivity matrix. We develop computationally efficient methods to perform CSS using a matrix congruence transformation and evaluate a range of different smoothing kernel choices on CSS performance. We find that smoothing substantially improves the identifiability, sensitivity, and test-retest reliability of high-resolution connectivity maps, though at a cost of increasing storage burden. For atlas-based connectomes (i.e. low-resolution connectivity maps), we show that CSS marginally improves the statistical power to detect associations between connectivity and cognitive performance, particularly for connectomes mapped using probabilistic tractography. CSS was also found to enable more reliable statistical inference compared to connectomes without any smoothing. We provide recommendations on optimal smoothing kernel parameters for connectomes mapped using both deterministic and probabilistic tractography. We conclude that spatial smoothing is particularly important for the reliability of high-resolution connectomes, but can also provide benefits at lower parcellation resolutions. We hope that our work enables computationally efficient integration of spatial smoothing into established structural connectome mapping pipelines.

**Highlights:**

- We establish a network equivalent of image smoothing for structural connectomes.
- Connectome Spatial Smoothing (CSS) improves connectome test-retest reliability, identifiability and sensitivity.
- CSS also facilitates reliable inference and improves power to detect statistical associations.
- Both high-resolution and atlas-based connectomes can benefit from CSS.

## 1. Introduction

Spatial smoothing is widely recognized as a crucial preprocessing step in many neuroimaging pipelines. It can increase the signal-to-noise ratio (SNR) by eliminating the high-frequency spatial components of noise [1–5] and is typically used in different neuroimgaing modalities such as structural magnetic resonance imaging (MRI) [6–8], functional MRI [9–13], positron emission tomography (PET) [14– 17], magnetoencephalography (MEG) [18, 19], electroencephalography (EEG) [20], and functional near-infrared spectroscopy (fNIRS) [21, 22]. As a result, options for spatial smoothing are provided in many neuroimaging toolboxes, such as AFNI [23], FreeSurfer [24], FSL [25], and SPM [26].

Structural connectivity computed from diffusion MRI tractography can be used to construct structural connectomes [27–29], and there is considerable interest in performing statistical inference on this graph representation of the brain [30, 31]. However, unlike image-based statistical inference, such data are currently not explicitly smoothed. Most structural connectomes are studied at the resolution of large-scale brain atlases comprising tens to hundreds of regions. The process of assigning tractography streamlines to such large-scale regions manipulates the data in a manner somewhat akin to smoothing. Nonetheless, the potential impact of additional (explicit) smoothing has not yet been evaluated. Moreover, given that connectomes are spatially embedded graphs, conventional univariate smoothing methods are not directly applicable to connectomes, and so smoothing methods tailored to connectome data are required.

High-resolution connectomes are a subset of connectomes that investigate the connectivity structure of the brain at the resolution of cortical vertices/voxels [32]. Recent studies highlight the advantages of investigating structural connectomes at this higher spatial resolution than atlases with coarse parcellations [32–39]. For example, high-resolution structural connectivity maps robustly capture intricate local modular structures in brain networks and provide insightful connectome biomarkers of neural abnormalities [36, 37, 40]. We recently established a computationally efficient framework to map high-resolution structural connectomes, and found that these connectomes enabled accurate prediction of individual behaviors and neural fingerprinting [32]. As part of this recent work, we implemented a preliminary method for connectome smoothing, building on earlier structural connectome smoothing approaches [33].

In this study, we extend our earlier work by formalizing the principles of Connectome Spatial Smoothing (CSS), aiming to develop efficient computational methods to facilitate connectome smoothing and determine optimal smoothing parameters. We investigate the impact of smoothing on high-resolution and atlas-based connectomes, quantifying its benefits for reliability, identifiability and statistical power. We anticipate that CSS will become a common step in connectome mapping workflows.

## 2. Materials and methods

### 2.1. Connectome Spatial Smoothing

Here, we develop an efficient and scalable method to enable smoothing of spatially-embedded high-resolution connectivity matrices. Unlike conventional spatial smoothing algorithms that are defined in terms of a single smoothing kernel, CSS is inherently bivariate and involves a pair of spatially distant smoothing kernels operating at the two ends of each connection. The framework developed here extends our recent work on high-resolution connectomes, where we first investigated the concept of connectome smoothing with a single kernel matrix [32]. We also acknowledge the seminal work of Besson and colleagues, who found that connectome smoothing improved the reliability of high-resolution connectomes [33], and other approaches in mapping continuous high-resolution connectomes with an implicit form of connectivity spatial smoothing [38, 39].

We use *A* to denote the symmetric connectivity matrix inferred from tractography, with size *v* × *v* where *v* is the total number of network nodes and element *A*(*i, j*) stores the streamline count between nodes *v*_*i*_ and *v*_*j*_. This matrix can be decomposed into two half-incidence matrices *U* and *V*, each of size *v* × *n*, where *n* is the total number of streamlines. These matrices encode the connectivity endpoint information, such that the streamline endpoint pairs are mapped to the columns of *U* and *V*. For instance, if the *k*th streamline ends in nodes *v*_*i*_ and *v*_*j*_, then the *k*th columns of *U* and *V* are vectors with a single non-zero element, with weight 1, respectively located at *U* (*i, k*) and *V* (*j, k*). This signifies that streamline *k* connects the endpoints *v*_*i*_ and *v*_*j*_. Fig. 1A-C demonstrates the decomposition of streamlines encoded in a connectivity matrix and the half-incidence matrix representations. Mathematically, the symmetric connectivity matrix is decomposed as follows:

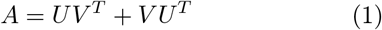

**Fig. 1.**
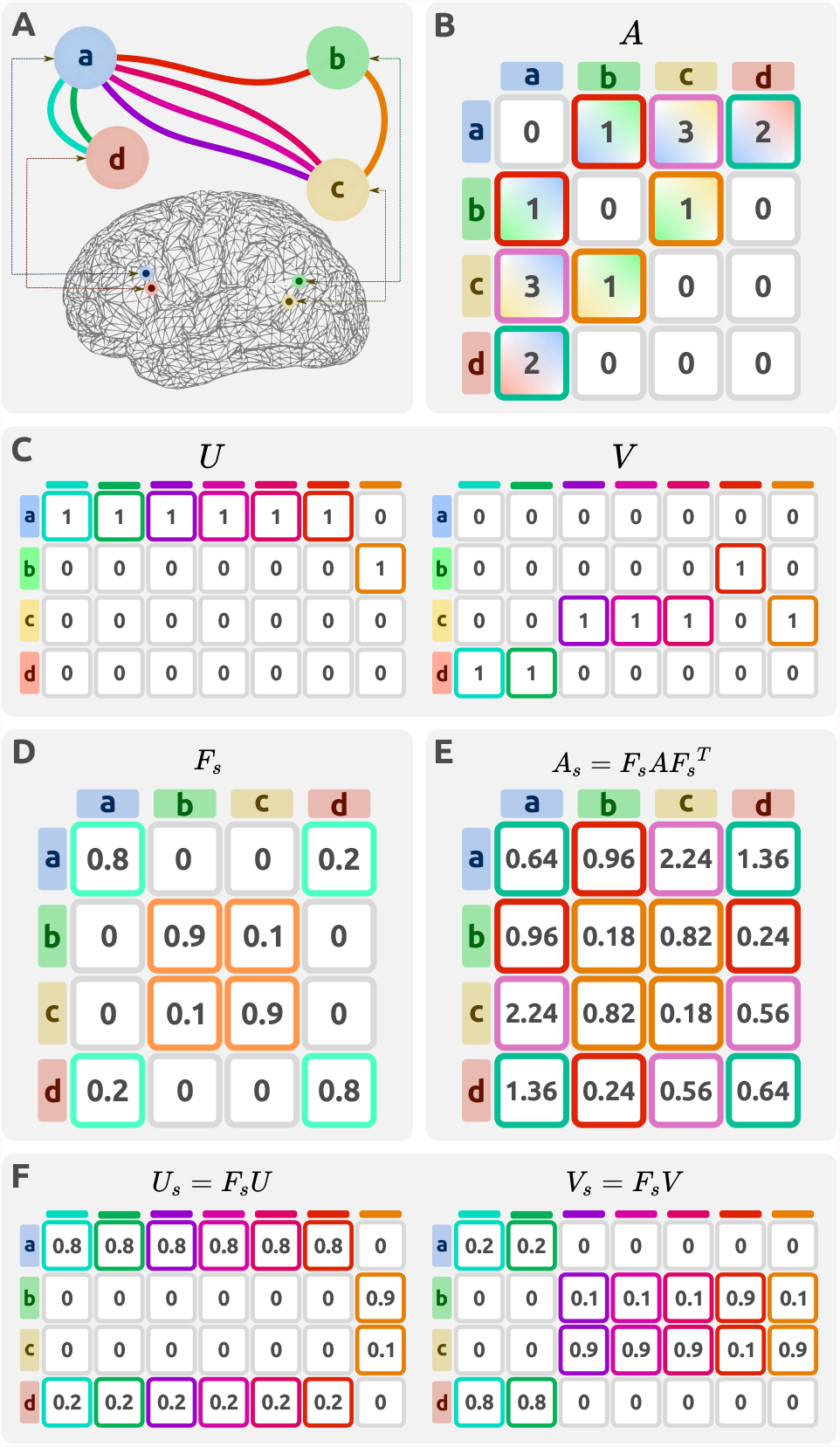
Illustrative example demonstrating the decomposition of streamlines into connectivity and incidence matrices. (A) A hypothetical network in which 7 streamlines connect four brain regions/nodes. The nodes (a,d) and (b,c) are selected to be spatially proximal. (B) Matrix *A* encodes the network in a 4×4 connectivity matrix. (C) The network can be alternatively represented by two half-incidence matrices *U* and *V*. (D) A connectome smoothing kernel *F*_*s*_ can be defined based on the pairwise geodesic distances between nodes. (E,F) The network representations can be spatially smoothed using CSS to produce a smoothed connectivity matrix *A*_*s*_ or a pair of smoothed half incidence matrices *U*_*s*_, *V*_*s*_.

Since the columns of the half-incidence matrices each represent a single endpoint associated with a spatial coordinate, a conventional spatial smoothing kernel can be applied to those columns, resulting in a pair of smoothed half-incidence matrices *U*_*s*_ and *V*_*s*_. As previously derived [32], a smoothed connectivity matrix can be constructed by combining the smoothed half-incidence matrices as follows:

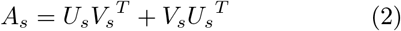

Here, we propose a simplification of this formulation, which leads to improved computational tractability. Let *F*_*s*_ denote a spatial smoothing kernel of size *v* ×*v*, such that column *i* of *F*_*s*_ stores the weights for a smoothing kernel spatially centered at the *i*th node of the network. This smoothing kernel can be used to compute the smoothed half-incidences:

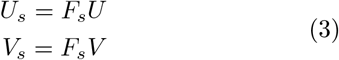

Smoothing can thus be represented as a linear transformation of each half-incidence matrix. Under this simplification, CSS reduces to a matrix congruence between the smoothed and initial connectivity matrices, which can be efficiently computed without using half-incidence matrices. Specifically, we have that:

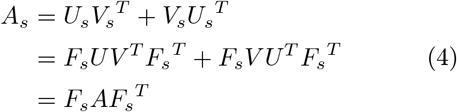

This equation shows that the smoothed connectivity *A*_*s*_ is a congruent transformation of the initial connectivity matrix *A*. Fig. 1D-F illustrates a simple example of this transformation. This simplification improves the computational feasibility since performing CSS is no longer dependent on the number of streamlines *n* which is typically greater than the number of non-zero connectome edges. The precise derivation of smoothing kernel matrix *F*_*s*_ is presented later in Section 2.6. Smoothing parameters.

### 2.2. Study design

We investigated the impact of CSS on connectomes mapped at different node resolutions. As detailed below, high-resolution (∼ 60k nodes) and atlas-based (∼ 300 nodes) connectomes were mapped for individuals using diffusion MRI and established whole-brain tractography methods. Data from two diffusion MRI acquisitions for each individual were used, enabling evaluation of test-retest reliability and identifiability across different smoothing parameters. Fig. 2 provides a brief overview of the study design. We next describe the diffusion MRI acquisition, whole-brain tractography and connectome mapping procedures, smoothing parameters, and the evaluation methodology used in this study.

**Fig. 2.**
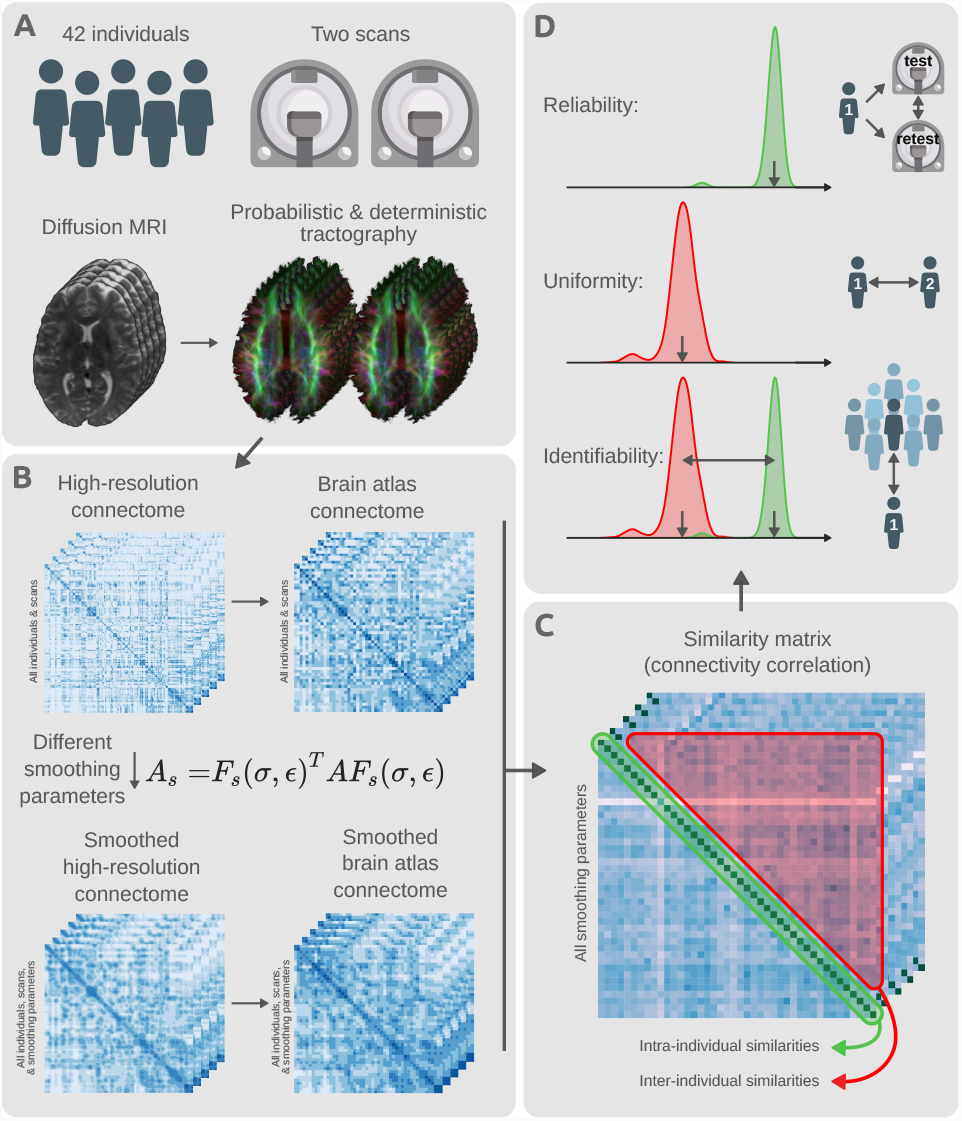
Schema of study design and methodology. (A) Test-retest diffusion MRI scans of 42 individuals were sourced from the Human Connectome Project. This provided a duplicate scan of every individual. Probabilistic and deterministic tractography were utilized to estimate whole-brain white matter fiber trajectories for all individuals and scans. (B) Tractography results were used to map unsmoothed structural connectomes using the high-resolution fsLR-32k surface mesh. Different smoothing parameters were used to transform the unsmoothed high-resolution connectomes into various CSS smoothed alternatives. All variants of smoothed and unsmoothed connectomes were also downsampled to connectivity maps on the HCP-MMP1.0 brain atlas comprising 360 cortical regions [48]. (C) All mapped connectomes were used to evaluate the level of similarity between connectivity maps of different scans (test and retest) for each combination of parcellation resolution and set of smoothing parameters. Both intra- and inter-individual similarities were computed for all pairs of scans. (D) The computed similarities were used to evaluate the level of connectome reliability, uniformity, and identifiability: reliability quantifies the average similarity of connectomes belonging to scans of the same individual; uniformity quantifies the average conformity of connectomes belonging to different individuals; identifiability measures the extent to which scans of the same individuals are differentiable from the rest of the group.

### 2.3. Imaging data acquisition and preprocessing

Imaging data were sourced from the Human Connectome Project (HCP) [41, 42]. We obtained the diffusion and structural MRI images from the 42 healthy young adults comprising the HCP test-retest cohort. For these individuals, two separate imaging sessions were conducted across two different days, with the intervening period between the test and retest scans ranging from 18 to 343 days. These duplicate individual scans enabled the assessment of both intra- and inter-individual variations in the mapped connectivity information. Diffusion MRI data were acquired using a 2D spin-echo single-shot multiband EPI sequence with a multi-band factor of 3 and monopolar diffusion sensitization. The diffusion data consisted of three shells (b-values: 1000, 2000, 3000 *s/mm*^2^) and 270 diffusion directions equally distributed within the shells, and 18 b=0 volumes, with an isotropic spatial resolution of 1.25mm [43]. We analyzed preprocessed diffusion data, where preprocessing was completed by the HCP team, using an established minimal preprocessing pipeline (v3.19.0). This included b=0 intensity normalization across scanning sessions, EPI and eddy-current-induced distortion corrections, motion correction, gradient nonlinearity correction, registration to native structural space, and masking the final data with a brain mask [44].

### 2.4. Connectome resolution

We mapped both high-resolution and atlas-based connectomes to evaluate the impact of CSS on different parcellation granularities. All high-resolution connectomes were mapped on the fsLR-32k standard surface mesh, comprising 32,492 vertices on each hemisphere [45]. This space is recommended for high-resolution cross-subject studies of diffusion MRI as it provides an accurate representation of the cortical surface with fewer vertices than the native mesh [44]. The combined left and right cortical surfaces consisted of 59,412 vertices after exclusion of the medial wall. This study used surface vertices as network nodes for the high-resolution connectome, consistent with many previous high-resolution connectivity approaches [32, 34, 35, 39, 46, 47]; since alternative approaches have instead mapped connectomes on the faces of the surfaces [33, 36, 37], we explain their duality as well as how CSS can be applied to both implementations in Supplementary Information Section S.1. CSS for connectivity mapped on vertices vs. faces. The high-resolution maps were downsampled to a lower spatial resolution defined by the HCP-MMP1.0 atlas comprising 360 cortical regions [48]. The downsampling procedure is detailed in the Section 2.7. CSS for atlas-based connectivity. In brief, the high-resolution connectivity matrix was aggregated across all vertices belonging to each atlas region such that the connectivity weight between two atlas nodes was equal to the sum of the connectivity weights over all high-resolution vertices connecting those atlas nodes. The subcortex was not included in either high-resolution or atlas-based connectomes.

### 2.5. Tractography and connectivity mapping

The impact of CSS was evaluated on both probabilistic and deterministic tractography algorithms. MR-trix3 software was used to perform tractography [49]. An unsupervised method was used to estimate the white-matter (WM), grey-matter (GM), and cerebrospinal fluid (CSF) response functions [50] for spherical deconvolution [51]. The fiber orientation distribution (FOD) in each voxel was estimated using a MultiShell, Multi-Tissue Constrained (MSMT) spherical deconvolution, which improves tractography at tissue interfaces [52]. The fsLR-32k surface mesh was used to generate a binary voxel mask at the interface between WM and cortical GM, from within which tractography streamlines were uniformly seeded at random coordinates from within this ribbon. Probabilistic tractography was performed by 2nd-order integration over fiber orientation distributions (iFOD2) [53]. Deterministic tractography was performed using a deterministic algorithm that utilized the estimated FOD with a Newton optimization approach to locate the orientation of the nearest FOD amplitude peak from the streamline tangent orientation (“SD_Stream”) [54]. Five million streamlines were generated for each tractography method for each scan.

A streamlines propagation mask was generated using the intersection of voxels with non-zero white matter partial volume as estimated by FSL FAST [55] and voxels with non-zero sub-cortical grey matter volume as estimated by FSL FIRST [56]. The sub-cortical GM was included in the propagation mask to preserve long streamlines relaying through the sub-cortex, only terminating streamlines at the boundaries of cortical GM or CSF. The streamline endpoints were then mapped to the closest vertex of the individual’s WM surface mesh (fsLR-32k) according to the Euclidean distance metric (see Supplementary Information Section S.2. Considerations for endpoint assignment for further detail). Streamlines ending far from the cortical vertices (>2mm) were discarded. The remaining streamlines were used to generate a 59,412 × 59,412 high-resolution connectivity matrix for each of the two sessions for each individual. These data form the input for evaluation of CSS as described in the following sub-sections.

### 2.6. Smoothing parameters

The matrix of spatial smoothing kernels, *F*_*s*_, determines the spatial distribution of smoothing weights. We use a Gaussian function to define kernel weights, *G*(*δ*), as a function of distance from the kernel center, *δ*, as given by:

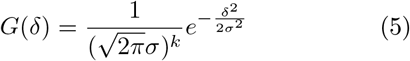

Where *k* is the dimension of the spatial kernel. The parameter *σ* is the standard deviation of the Gaussian distribution which determines the strength of smoothing. In this study, smoothing was applied to the cortical surface mesh (*k* = 2) and was quantified by the geodesic distance over the surface mesh. This geodesic distance metric indicates the shortest spatial path between two points that is constrained to lie entirely on the surface mesh [57]. Despite each subject possessing the same set of vertices, the smoothing kernel was computed separately for each scan, based on the precise inter-vertex geodesic distances on the white-matter surface mesh of each individual scan.

To compare the impact of different kernel standard deviations, smoothing kernels were computed with 1, 2, 3, 4, 6, 8, and 10mm FWHM (full width at half maximum) 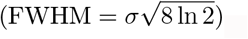.

A second parameter that can impact smoothing is *truncation* of the kernel. As the Gaussian distribution decays exponentially with distance, the kernel is effectively zero for sufficiently large distances, and so contributions can be ignored with minimal loss of precision. Truncation results in a sparse smoothing kernel, enabling computationally efficient smoothing of high-resolution connectomes. Here we studied the effect of the truncation threshold, *ε*, which is defined as the fraction of the kernel integral discarded as a result of kernel truncation (Fig. 3A): for each value of FWHM, we generated three kernels for assessment, corresponding to *ε* = {0.1, 0.01, 0.001}. This truncation can alternatively be expressed as a kernel radius *R* (which has benefits both conceptually and programmatically):

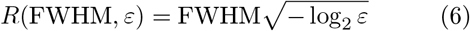

Proof of this relationship is provided in the Supplementary Information Section S.3. Thresholding radius.

**Fig. 3.**
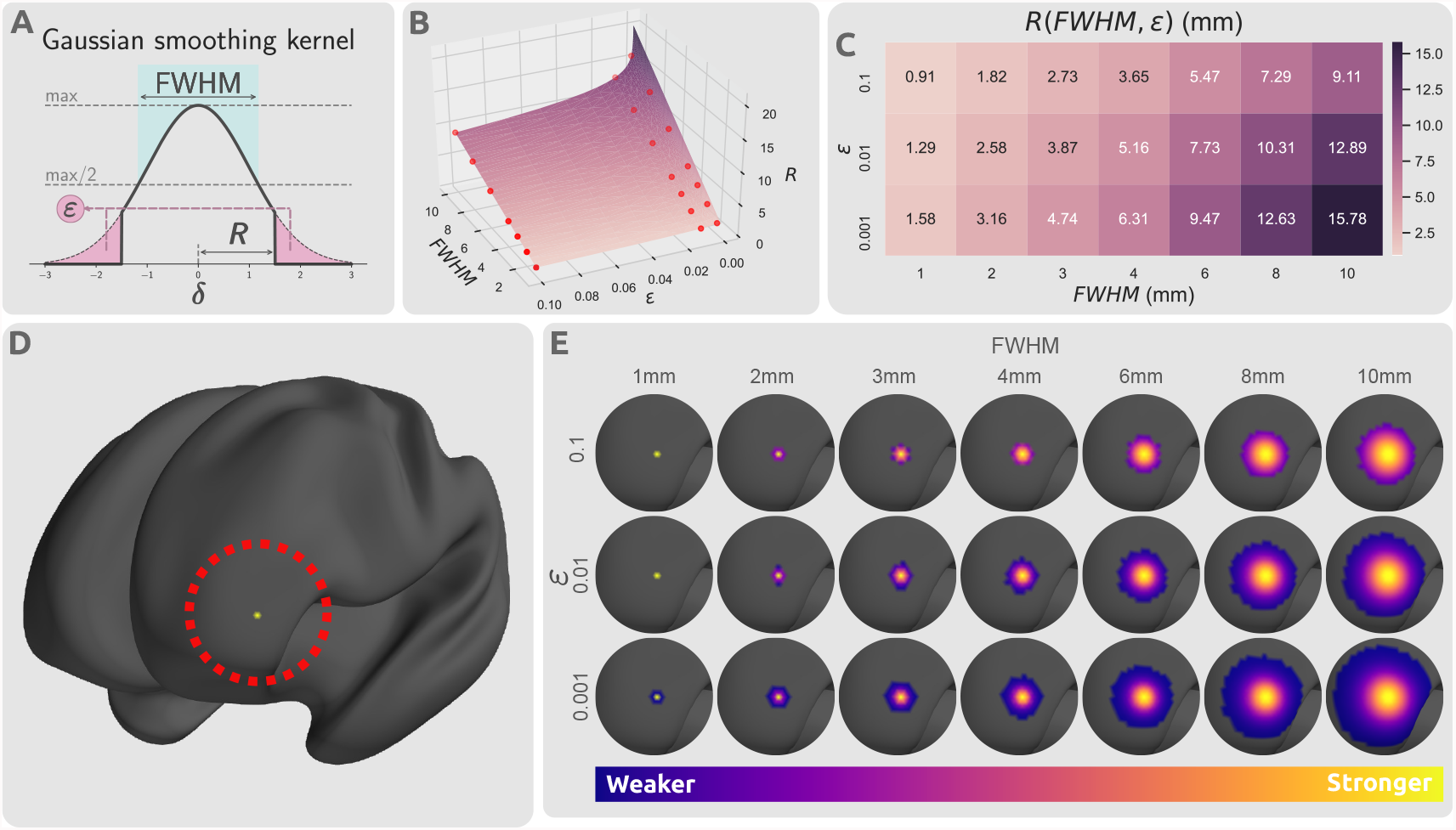
Impact of kernel parameters on truncated kernels. (A) Distribution of a truncated Gaussian kernel with smoothing parameters FWHM, *ε*, and *R*(FWHM, *ε*). FWHM determines the standard deviation of the Gaussian kernel, *ε* dictates the proportion of the kernel that is truncated, and *R* determines the threshold radius beyond which the kernel is set to zero. The thick black line represents the smoothing kernel as a function of the smoothing parameters. (B) Smoothing parameter space of FWHM, *ε*, and *R*. The parameter space plane shows the value of smoothing kernel radius *R* as a function of kernel standard deviation, FWHM, and truncation threshold, *ε*. The radius is linearly related with FWHM, but log-linearly with the inverse of *ε*. The red points indicate the selected smoothing parameters from the parameter space that were used in this study. (C) The values of kernel truncation radius at the respective smoothing parameters selected for FWHM and *ε*. (D) A sample cortical vertex in the left frontal lobe of an inflated cortical mesh. (E) The respective column of the smoothing kernel *F*_*s*_ for the vertex shown in panel (D) with different smoothing parameter choices projected on the cortical surface.

Fig. 3A shows the influence of FWHM and *ε* on the truncated kernel. Fig. 3B,C show the relationship between standard deviation, truncation threshold and radius. Truncated kernels were generated with nonzero kernel weights only at locations with distance less than *R*(FWHM, *ε*) from the kernel center. Consequently, kernels were re-normalized such that for every vertex the column sum of *F*_*s*_ was 1.0 despite truncation. This step removes any artifacts introduced by mesh vertex density variations (such as amplification of signal in regions with high vertex density). Fig. 3D,E demonstrate the spatial distribution of a single row of this smoothing kernel over a sample cortical surface mesh.

### 2.7. CSS for atlas-based connectivity

As described in Section 2.4. Connectome resolution, smoothed versions of the parcellation-based atlas-resolution connectome can be computed by first applying smoothing to the high-resolution connectome, then aggregating the connectivity values within the vertices corresponding to each atlas parcel. This approach however necessitates the high storage and computational complexity demands of high-resolution connectome data. We therefore derived a more computationally efficient procedure to perform CSS on atlas-based connectomes.

A brain parcellation atlas can be denoted by a binary *p* × *v* matrix *P*, where *p* is the number of brain regions in the atlas, such that the *i*th row of *P* is a binary mask of vertices belonging to the *i*th atlas region and each vertex belongs to at most one region (a “hard parcellation”). An atlas-based connectivity map *A*_*p*_ can be represented by the matrix multiplication *A*_*p*_ = *PAP* ^*T*^ : this operation reduces the *v* × *v* high-resolution connectivity *A* to a *p*× *p* atlas connectivity map *A*_*p*_. To smooth *A*_*p*_, the high-resolution connectivity matrix *A* can be smoothed to *A*_*s*_ and then downsampled to create the smoothed atlas connectivity map *A*_*sp*_. An equivalent approach is to first spatially smooth every row of the brain atlas *P*, and then normalize every column to produce a smoothed “soft parcellation” *P*_*s*_ = *PF*_*s*_, where each region is now defined as a weighted probability map across vertices and vertices can have non-zero membership to multiple regions. This enables direct computation of smoothed parcellation-based connectome matrix *A*_*sp*_ without necessitating computation of the smoothed high-resolution connectome matrix *A*_*s*_ (see Supplementary Information Section S.4. CSS for atlas-based connectivity for detail):

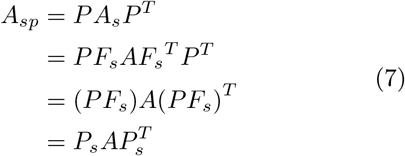

### 2.8. Connectome similarity

To evaluate the potential advantages of smoothing, a measure of similarity based on Pearson’s correlation was used to quantify the conformity of two connectivity maps [32, 58]. To compute the similarity between two networks *A*_1_ and *A*_2_, first, Pearson’s correlation was computed for all respective rows of the connectivity matrices, yielding *v* correlation coefficients, each indicating the connectivity similarity of a single node; these correlations were then averaged over all nodes to produce a single value indicating the similarity of two connectomes. This measure was used to quantify both intra- and inter-individual connectome matrix similarities.

### 2.9. Evaluation metrics

Direct connectome comparisons were performed within each combination of: tractography algorithm (deterministic and probabilistic); parcellation resolution; and network smoothing parameters. Within each of these configurations, smoothed structural connectomes were generated independently for the two scanning sessions for each of 42 participants. For each scan in session 1, its similarity to every session 2 scan (1 intra-individual and 41 inter-individual) was computed; aggregated across all individuals, this process yielded 42 values comparing connectomes of the same individual (intra-individual similarities), and 42× 41 measuring the similarity between connectomes of different individuals (inter-individual similarities). The intra-individual similarities were averaged to form a measure of connectome *reliability μ*_*intra*_, indicating the extent of consistency of mapped connectomes for an individual; similarly, the inter-individual similarities were averaged to yield a measure of population uniformity of the connectivity maps *μ*_*inter*_. Ideally, connectomes should be reliable (i.e. high *μ*_*intra*_) and preserve inter-individual differences (i.e. low *μ*_*inter*_). Hence, high reliability and low population uniformity is desirable.

To evaluate the extent to which an individual’s connectome is unique, we adopted an established identifiability framework [59]. *Identifiability* quantifies the extent to which an individual can be differentiated from a larger group based on a set of individual attributes. Here, identifiability was measured by the effect size of the difference in the means of intra-individual and inter-individual similarities [32]:

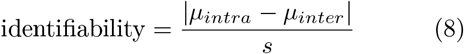

Where *μ*_*intra*_ and *μ*_*inter*_ are the mean of the two intra- and inter-individual similarity distributions and *s* is the pooled standard deviation of the two distributions.

### 2.10. Evaluating statistical power with atlas-resolution smoothing

Generally, smoothing can result in a loss of effective spatial resolution, blurring, and shifting or merging of adjacent signal peaks [60–63], but is necessary to strike a compromise between sensitivity and specificity [64]. Hence, we investigated the impact of CSS on mass univariate significance testing of associations between cognitive performance and atlas-based structural connectivity. Given that structural connectivity and cognition are known to be associated [32], we tested whether the use of CSS would improve power to detect such associations. For each pair of regions in the parcellation atlas, Pearson’s correlation coefficient was used to test for an association between connectivity strength and a previously established measure of overall cognitive performance [65]; This cognitive measure was generated from a data-driven behavioral dimension derived from independent component analysis (ICA) of 109 behavioral items. This yielded a correlation coefficient for each pair of regions. Age and sex were regressed out from the cognitive measure as confounds. This was repeated across 100 bootstrap tests each including 90% of the sample (N=35) to increase the robustness of the comparisons against individual effects.

To generate a distribution of correlation coefficients under the null hypothesis of an absence of association between connectivity and cognitive performance, we randomized cognitive scores between individuals and recomputed all correlation coefficients; this was repeated for 1000 randomizations (10 randomizations within each bootstrap sample), yielding 1000 correlation coefficients representing the null distribution for each connection. For a range of correlation coefficient thresholds from 0.1 to 0.5 (which indicate small to large effects according to Cohen’s conventions [66]), we counted the number of suprathreshold connections in both the empirical and randomized data (averaged across the 1,000 randomizations).

From these data, we generated an ROC (Receiver Operator Characteristic) curve as follows. The number of suprathreshold connections in the empirical data was assumed to give the combined number of true positives (*TP*) and false positives (*FP*), while the average total number of suprathreshold connections in the randomized data estimated the total number of *FP* (Fig. 4A). The combined number of false negatives (*FN*) and true negatives (*TN*) was determined by subtracting *TP* +*FP* from the total number of connections. Finally, since the true underlying effect was unknown, we assumed a false omission rate of 1%, i.e., 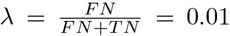. This assumption enabled estimation of sensitivity 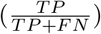 and specificity 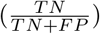 that were used to generate the ROC curve. We ensured that our estimates were robust to the choice of *λ* (see Supplementary Information Section S.5. Replication of ROC curve estimates for detail). This process was repeated independently for various smoothing kernels, and for data generated using both deterministic and probabilistic tractography algorithms, to investigate the impact of CSS on the statistical power to detect associations between cognitive performance and connectivity.

**Fig. 4.**
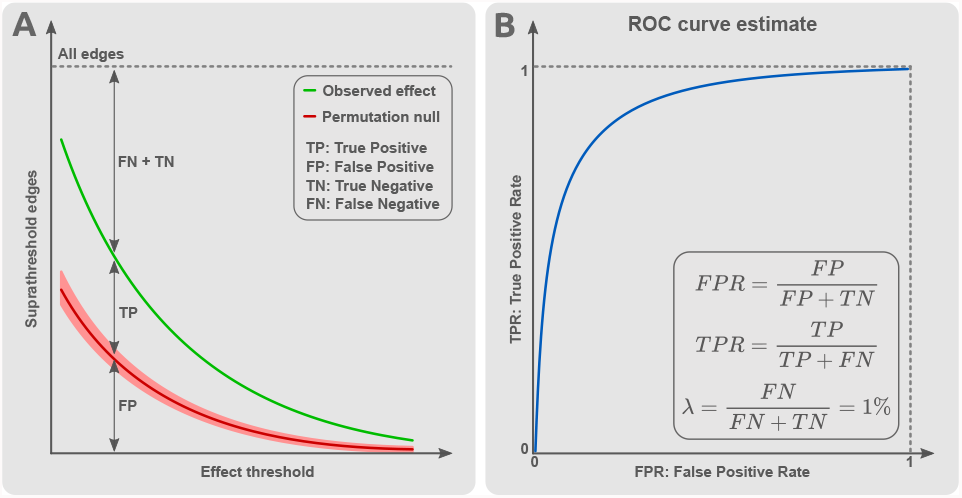
Estimation of receiver operating characteristic (ROC) curves for mass univariate testing of associations between cognitive performance and structural connectivity. For each pair of regions in the parcellation atlas, Pearson’s correlation coefficient was used to test for an association between connectivity strength and a previously established measure of overall cognitive performance. (A) For a given effect size threshold (horizontal axis), the number of suprathreshold connections (vertical axis) yielded the combined number of true positives (*TP*) and false positives (*FP*), indicated by the green line. The red line indicates the total number of *FP*, determined by randomizing cognitive scores between individuals and recomputing all correlation coefficients (1000 randomizations; mean & 95% confidence interval shown). (B) Assuming a constant value for the false omission rate 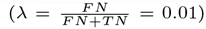, an ROC curve can be estimated for different effect size (correlation coefficient) thresholds. TPR: true positive rate. FPR: false positive rate.

Additionally, we tested the replicability of the suprathreshold effects in a test-retest comparison to evaluate the replicability of the observations before and after smoothing. At each utilized threshold value, for every edge that was suprathreshold in the data from either session 1 or session 2, we calculated the difference in correlation coefficient between the two sessions. This provided a distribution of effect differences observed across a range of effect thresholds. Thus, a lower average effect difference indicated higher consistency of the connectivity-behavior observations and higher replicability of the findings.

## 3. Results

We investigated the utility of CSS for high-resolution and atlas-based connectomes, focusing on connectome reliability and identifiability as well as computational and storage requirements. We recommend optimal smoothing kernels for connectomes mapped with deterministic and probabilistic tractography, and we demonstrate that smoothing improves the statistical power to detect associations between connectivity and cognitive performance.

### 3.1. High-resolution connectome storage size

High-resolution connectomes require considerable storage and computational resources, and CSS can increase this burden, due to reductions in matrix sparsity. Fig. 5 summarizes the sizes of stored connectomes for various kernels. Kernels with larger FWHM and/or more lenient truncation thresholds incur greater storage demands for high-resolution connectomes. We found that the kernel radius *R*(FWHM, *ε*), which is dependent on both parameters, was a reasonable predictor of connectome size. We also observed that connectomes mapped using probabilistic tractography were approximately an order of magnitude larger than their deterministic counterparts both prior to smoothing (∼ 10MB for probabilistic and ∼ 1MB for deterministic) and after performing CSS with identical smoothing parameters.

**Fig. 5.**
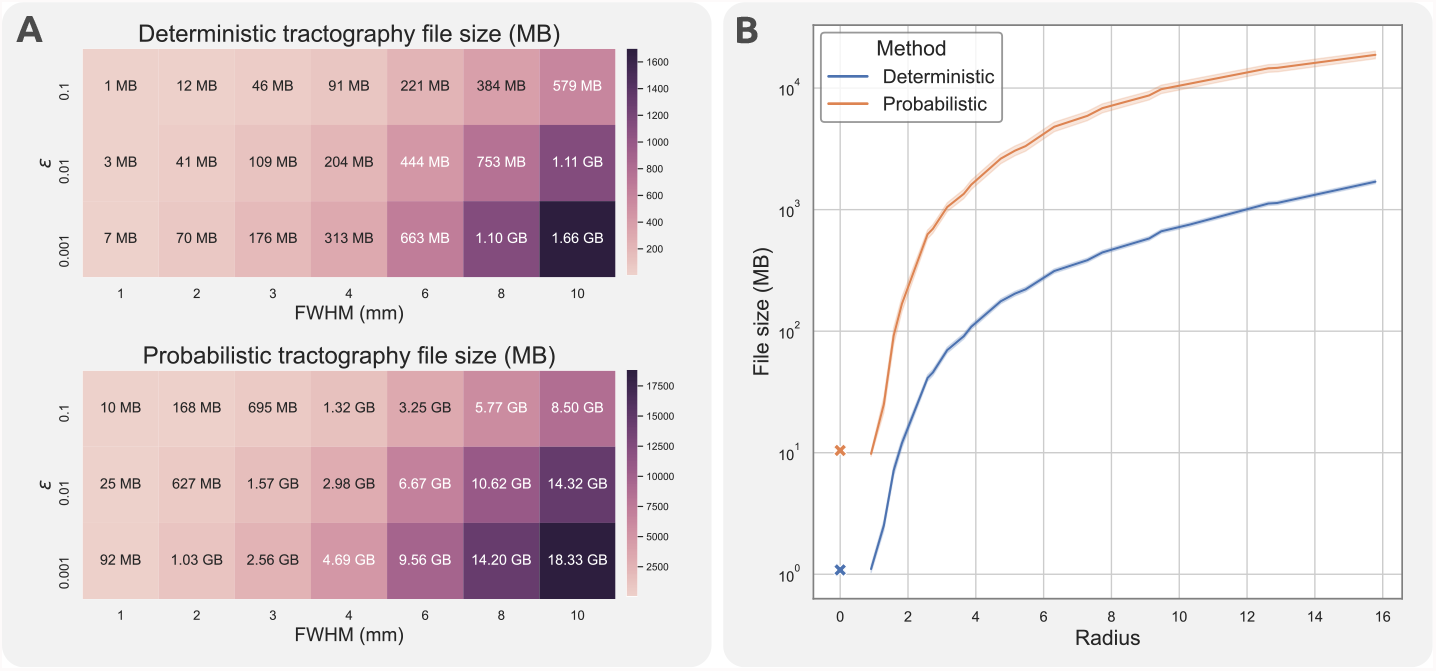
Impact of CSS on connectome storage requirements. (A) Tables show the mean storage size of individual connectomes mapped using deterministic (upper) and probabilistic (lower) tractography following smoothing, as a function of truncation threshold and full-width at half maximum (FWHM) of smoothing kernel (B) The relationship between the kernel radius and file size of individual connectomes. Results for connectomes with no smoothing are marked with an x. File sizes are plotted using a logarithmic scale. Shaded bands indicate one standard deviation from the mean.

### 3.2. Identifiability and reliability

Fig. 6 summarizes the impact of CSS on the identifiability and reliability of high-resolution structural connectivity maps. We observed that both larger FWHM values and smaller truncation thresholds (i.e., larger *R* in both cases) consistently improved connectome reliability (mean intra-subject similarity). While high-resolution connectomes without smoothing had a relatively low reliability (*μ*_*intra*_ < 0.2), CSS with kernels as little as 3-4mm FWHM resulted in a substantial increase in reliability (*μ*_*intra*_ > 0.5), with reliability exceeding 90% (*μ*_*intra*_ > 0.9) achieved in some scenarios.

**Fig. 6.**
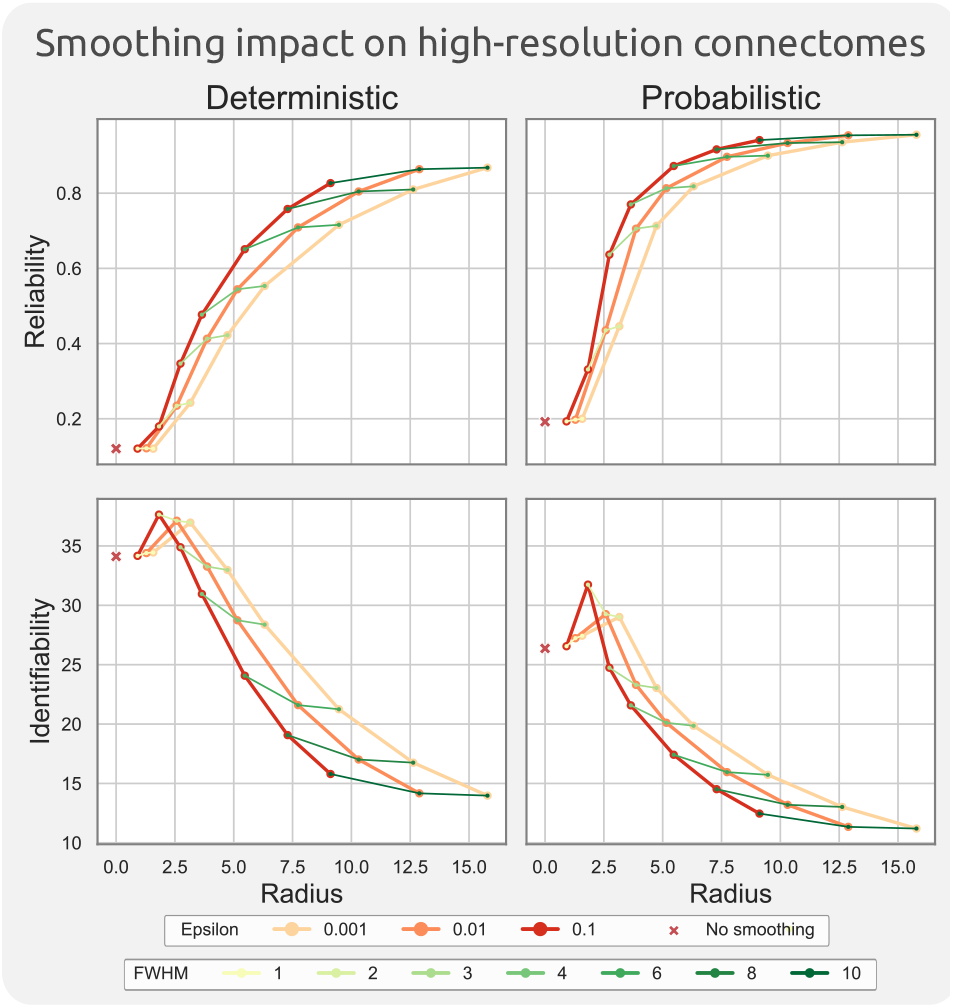
Impact of CSS on high-resolution connectomes for a range of different kernel parameters. Reliability (first row) and identifiability (second row) are reported for deterministic (left column) and probabilistic (right column) structural connectomes mapped at the resolution of cortical vertices. Results for connectomes with no smoothing are marked with an x in each plot. Kernel truncation thresholds, *ε*, are colored using warm colors such that each line connects points with equal *ε*; similarly, FWHM is colored using shades of green.

CSS also impacted connectome identifiability. We observed that while CSS with a 2-4mm FWHM kernel improved the identifiability of connectomes, CSS with larger FWHM was detrimental for individual identifiability, such that CSS with a 10mm FWHM resulted in more than 50% reduction in identifiability. For both identifiability and reliability measures, CSS was more sensitive to a change in kernel FWHM in contrast to the truncation threshold *ε*. Increasing the truncation threshold from *ε* = 0.01 to *ε* = 0.001 had negligible impact on either measure.

Tractography algorithm choice also impacted reliability and identifiability. High-resolution connectomes mapped using deterministic tractography had relatively lower reliability (10-20% lower), but higher identifiability (20-30%), compared to their probabilistic counterparts with identical CSS parameters.

Fig. 7 shows the impacts of CSS with different kernel parameters on an atlas-parcellation-based structural connectome. In agreement with the high-resolution analyses, we observed both that increases in FHWM and decreases in kernel truncation thresholds led to improved connectome reliability, and that this improvement in reliability comes at the expense of reduced identifiability. Without CSS, the atlas-based connectomes were already relatively reliable (deterministic: 92%, probabilistic: 98%). Use of the largest smoothing kernel increased these to 97% and 99%, respectively, albeit at the cost of a small reduction in identifiability (from 7.8 to 7.2 for deterministic and from 6.8 to 6.1 for probabilistic). Changing kernel extent from *ε* = 0.01 to *ε* = 0.001 again had no considerable impact on reliability or identifiability. The magnitude of influence of CSS on the atlas-resolution connectomes was comparatively smaller than the effects observed at the higher resolution.

**Fig. 7.**
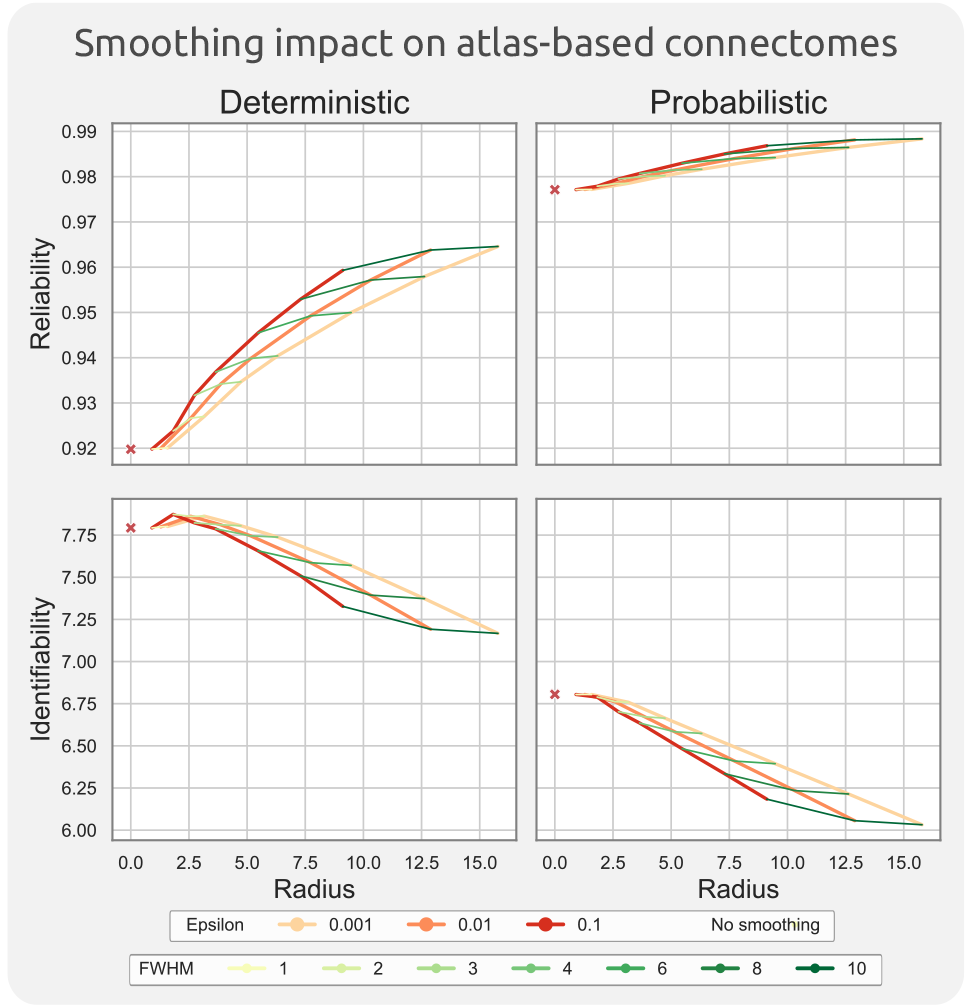
Impact of CSS on atlas-based connectomes, for a range of different kernel parameters. The connectome reliability (first row) and identifiability (second row) are reported for deterministic (left column) and probabilistic (right column) structural connectomes mapped at the resolution of atlas parcels. The unsmoothed atlas-based connectivity results are marked with x in each plot. Kernel truncation thresholds, *ε*, are colored using warm colors such that each line connects points with equal *ε*; similarly FWHM is colored using shades of green.

All in all, we observed that the advantages of CSS for high-resolution were maximized with 3-6mm FWHM kernels; larger smoothing kernels (>6mm FWHM) could deteriorate high-resolution identifiability for the sake of reliability. In contrast, identifiability of the atlas-resolution maps were less sensitive to larger smoothing kernels, and thus kernels of 6-10mm FWHM can be used to improve reliability with proportionally smaller losses in identifiability. To achieve similar reliability and identifiability, connectomes generated using deterministic tractography were found to require CSS with larger smoothing kernels compared to their probabilistic counterparts. Finally, it should be noted that the optimal CSS kernel parameters essentially depend on the application for which the connectome will be used.

#### 3.2.1. Case Study: Impact of CSS on statistical power

Finally, we investigated whether CSS can improve statistical power to detect associations between structural connectivity and cognitive performance. For this study we used FWHM = 8mm and *ε* = 0.01, based on the results reported above. For this case study, we considered the mapped atlas-based connectomes and computed Pearson’s correlation coefficient between streamline counts and cognitive performance for each pair of regions. ROC curves were then computed for each case, as described in the Methods, to determine whether CSS improved statistical power to identify associations between connectivity and cognitive performance.

First, we tested whether the magnitude of effect in the set of suprathreshold connections (i.e., connections with a correlation coefficient exceeding a fixed threshold) were replicable between the test and retest datasets. We found that CSS improved replicability in suprathreshold connections, particularly more so for connectomes mapped with probabilistic tractography (Fig. 8A); this suggests that CSS can improve the reproducibility of mass univariate testing on connectomes.

**Fig. 8.**
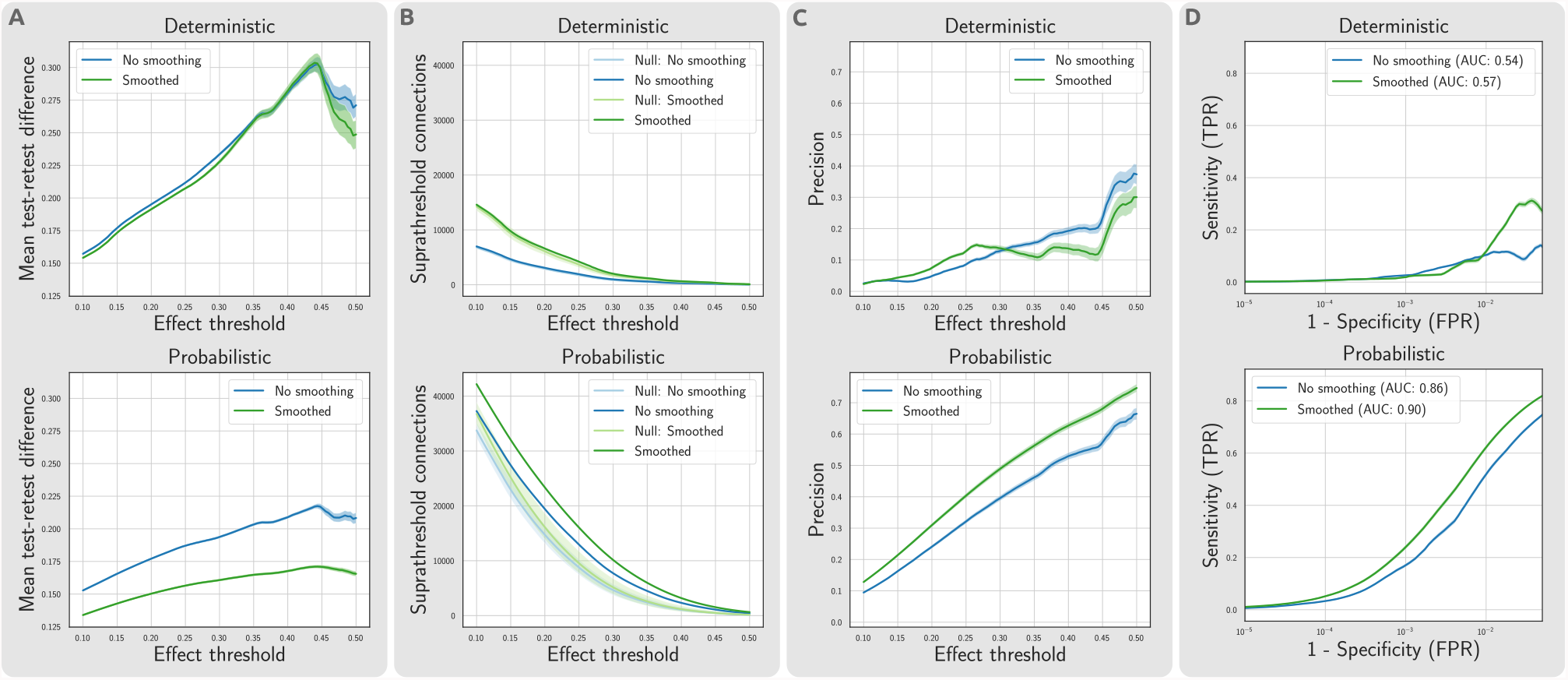
Impact of CSS on statistical power of mass univariate testing on atlas-based connectomes, based on an exemplar dataset examining correlations between structural connectivity and cognitive performance. (A) Replicability of suprathreshold connections between test and retest datasets: a lower difference of the observed effect magnitude between test and retest is favorable in terms of replicability. (B) The number of suprathreshold connections as a function of the effect size threshold was compared with a null distribution from permutation. To assess the predictive utility of the connectomes, precision, sensitivity, and specificity was estimated from a comparison with the null. (C) Precision was calculated from the ratio of supra-threshold edges found in empirical data compared to the null model at different effect thresholds. (D) ROC curves were estimated to demonstrate the respective changes in sensitivity 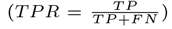 and specificity 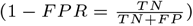 of the edges selected at different effect thresholds. The analyses were repeated across bootstrap samples to provide a robust estimate of statistical power. Shaded lines indicate 95% confidence intervals. The Area Under Curve (AUC) metric was reported to facilitate performance comparison. Abbreviations: TP: True Positive, FP: False Positive, TN: True Negative, FN: False Negative, TPR: True Positive Rate, FPR: False Positive Rate.

Next, we enumerated the number of suprathreshold connections as a function of the effect threshold (Fig. 8B). While the proportion of suprathreshold connections increases following smoothing for the empirical data, indicating a potential gain in sensitivity, a similar increase in the randomized (null distribution) data suggests that this may come at the expense of poorer specificity. For connectomes mapped with deterministic tractography, the numbers of suprathreshold connections for the empirical and randomized data are separated by a comparable gap, irrespective of whether CSS was performed. For probabilistic tractography, the number of suprathreshold connections for the randomized data was comparable with and without smoothing, whereas CSS resulted in a substantially greater proportion of suprathreshold connections for the empirical data. This suggests that CSS can improve the statistical power of mass univariate testing performed on connectomes mapped with probabilistic tractography, without a substantial loss in specificity.

To further investigate these effects, we considered precision 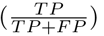 as a function of effect size threshold (Fig. 8C); and from this, generated ROC curves (Fig. 8D). Performing CSS on connectomes mapped from probabilistic tractography improves the precision and sensitivity of the inference. This improvement is also partially observed for connectomes mapped from deterministic tractography only for smaller effect thresholds (*r* < 0.3). Taken together, these results suggest that CSS is particularly beneficial to improving the statistical power of inference performed on connectomes mapped with probabilistic tractography; in contrast, for connectomes mapped with deterministic tractography, the benefit of CSS is marginal and possibly detrimental for larger effect size thresholds (*r* > 0.3). More importantly, CSS improved replicability with minimal impact on the statistical power for connectomes mapped with both tractography algorithms.

## 4. Discussion

In this study, we established a computationally efficient formalism for connectome smoothing and demonstrated that our Connectome Spatial Smoothing (CSS) method can benefit the analysis of atlas-based and high-resolution connectomes. Our results demonstrate that CSS impacts different aspects of connectivity mapping analyses, including individual reliability, inter-individual variability, and the inter-scan replicability of brain-behavior statistical associations, as well as computational storage demands. The choice of smoothing kernel parameters involves a trade-off between connectome sensitivity and specificity: larger kernels (higher FWHM and lower *ε*) improve connectome sensitivity, but are detrimental to connectome specificity. It is therefore important to select a level of smoothing that strikes a balance between these competing factors. In the following sections, we provide some guidelines for selecting optimal smoothing parameters and discuss the implications of performing CSS for connectome reliability, identifiability, storage requirements, and statistical power.

### 4.1. Appropriate smoothing parameters

Our results indicate that CSS differentially affects the characteristics of structural connectivity matrices mapped with different tractography methods and parcellation resolutions. Although we cannot suggest a one-size-fits-all smoothing kernel, our findings can guide selection of appropriate CSS smoothing kernels in future studies. Table 1 provides some rules of thumb for selecting a level of spatial smoothing which aims to achieve a balance between reliability and identifiability, while also considering storage demands. In general, high-resolution connectomes benefit from smaller FWHM compared to atlas-based connectomes, and deterministic maps require larger FWHM than their probabilistic counterparts to achieve the same level of reliability. However, the goals of the analysis at hand must be considered when selecting the level of smoothing. For example, if the goal is to identify an individual from a group based on their connectome, deterministic tractography and a smaller FWHM than recommended in Table 1 may be desirable. On the other hand, if one wishes to build a reliable consensus structural connectome that is robustly consistent across individuals, a higher FWHM than recommended in Table 1 may be favored. A value of 0.01 is suggested universally for the kernel truncation threshold *ε*, as smaller thresholds yield negligible impacts on identifiability and reliability whilst incurring much greater storage costs.

**Table 1.**
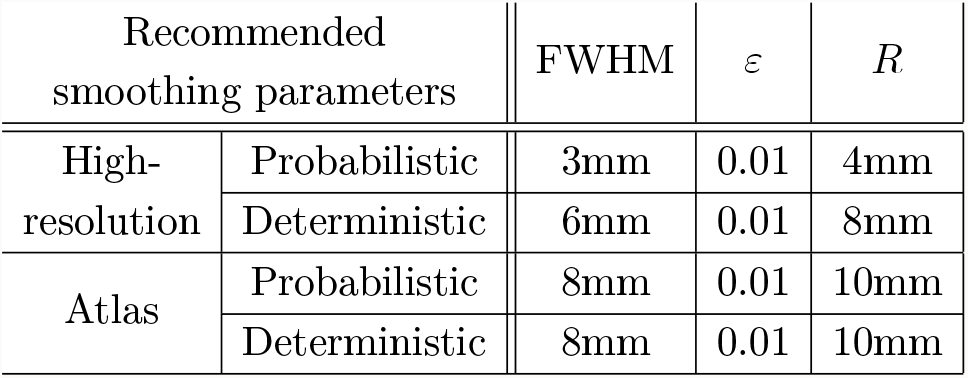
Recommended smoothing parameters. This table provides rule of thumb recommendations for CSS smoothing kernels of different variants of structural connectomes. In general, connectomes at the resolution of a brain atlas can benefit from larger CSS kernels compared to high-resolution connectomes. High-resolution connectomes computed from probabilistic tractography are advised to be smoothed less than their deterministic counterparts. Reducing epsilon below 0.01 is unfavorable and computationally costly. The rounded values for kernel radius *R*(FWHM, *ε*) provide sensible approximations.

Without CSS, connectomes mapped from deterministic tractography were found to yield higher identifiability; conversely, connectomes mapped from probabilistic tractography were more reliable. This is in line with previous reports suggesting that probabilistic tractography achieves higher sensitivity, lower specificity, and lower interindividual variability, compared to deterministic approaches [67–70]. Given that many factors other than reliability and identifiability would affect the choice of tractography algorithm, we suggest that CSS could be leveraged to achieve a balance between reliability and identifiability of the selected tractography algorithm. Hence, we could take advantage of a comparatively larger kernel for deterministic tractography approaches to match the reliability and identifiability of the probabilistic counterpart.

Our results highlight that CSS is a critical step to improving the reliability of high-resolution connectomes. High-resolution connectivity mapping is particularly sensitive to noise, artefacts, and registration misalignment, all of which can be alleviated—to a certain extent—with the new CSS formalism developed here.

### 4.2. Connectome reliability

Structural connectivity maps are commonly used in research to draw statistical inferences regarding associations between brain connectivity and different aspects of human cognition, behavior, and mental health [71–76]. The statistical power of such inferences can depend on the reliability of the measure under study: a connectivity measure that can be reliably assessed for all individuals can potentially improve the characterization of brain-behavior associations. However, improvements in reliability achieved by increasing the level of smoothing come at the expense of poorer spatial specificity and increases in connectome storage and computational requirements. CSS enables researchers to balance this trade-off to match the goals of the analysis at hand. Commonly used atlas-based connectivity maps are comparatively reliable, even without any smoothing, since the reduced spatial resolution of inter-subject correspondence imposed by a parcellation performs an operation comparable to smoothing. Nevertheless, we found that CSS could marginally improve the reliability of atlas-based connectomes.

### 4.3. Individual identifiability

The concept of neural fingerprinting has emerged in recent years which considers the challenge of identifying an individual from within a large group of others, based on their connectome or other neuroimaging data [58]. While the efficacy of a measure at individual identification does not necessitate existence of behavioral and pathological biomarkers in individuals, it could still be conceived as an indicator of the strength of such individual brain-behavior associations. By reducing the impact of noise and registration misalignments, CSS can enhance detection of individual differences in connectivity maps, enabling clearer differentiation of individuals and thus potentially improve the accuracy of neural fingerprinting. Our findings suggest that a minimal smoothing kernel of 2mm FWHM improves both reliability and identifiability of high-resolution connectivity matrices. Implementing CSS with larger kernels (i.e. > 2mm FWHM) further enhances connectome reliability substantially, but results in a gradual reduction in identifiability due to loss of individual identifiers by spatial blurring. Smoothing the high-resolution connectivity maps beyond 6mm FWHM is unnecessary because gains in reliability diminish, despite detrimental impacts on identifiability, spatial specificity, and storage requirements.

### 4.4. Storage requirements

It is important to consider the storage demands and associated computational burdens of handling smoothed connectome data. If the connectome size is larger than a gigabyte or so, handling the file (loading into memory and conducting analyses) can become unacceptably time-consuming. Even with the assistance of high-performance computing infrastructure, any benefits of using connectomes larger than a few gigabytes might not outweigh the time and resources required to process the larger files. This especially limits the extent of smoothing for connectomes generated using probabilistic tractography, which can grow to more than a few gigabytes when smoothed above 4-6mm FWHM. In contrast, connectomes mapped with deterministic tractography can be smoothed further whilst remaining highly computationally feasible. Nevertheless, if greater smoothing is essential in a study, a high-performance computing platform with access to adequate memory can be used to process smoothed connectomes (potentially without use of sparse matrix data structures), which may take tens of gigabytes of memory per individual connectome.

### 4.5. Implications on atlas resolution

Our results highlight the impact of CSS on structural connectomes mapped both at the high resolution of individual surface vertices, and the lower resolution of a brain atlas. While the findings vary in terms of magnitude of influence, a common pattern is visible across resolutions: higher FWHM results in a more reliable connectome, yet higher FWHM reduces the identifiability of connectomes. We developed computationally efficient methods to perform CSS at both resolutions. Atlas-based connectivity matrices have a relatively small memory footprint (<1MB), and thus they can be processed and stored efficiently, regardless of the level of smoothing. Similar to the high-resolution connectomes, when using an atlas parcellation, probabilistic and deterministic tractography approaches have complementary attributes when comparing reliability and identifiability: connectomes mapped from probabilistic tractography achieve better reliability compared to their deterministic counterparts, whereas deterministic connectomes can better reveal individual differences. This is one possible factor that can guide the choice between deterministic and probabilistic tractography algorithms. However, CSS can be used to increase the reliability of connectomes mapped from deterministic tractography to match the reliability of the probabilistic approach.

Finally, the atlas-based smoothing results suggest that probabilistic maps are to a certain extent representative of highly smoothed deterministic ones. In other words, more smoothed deterministic maps were analogous to less smoothed probabilistic maps, as the probabilistic evaluation curves in Fig. 7 seem to be a continuation of the deterministic curves. This observation is in agreement with prior expectations given the mechanisms used to generate the data, as probabilistic tractography-based connectivity has an intrinsic spatial smoothness due to the stochastic variability in streamline propagation. The proposed method to perform CSS on atlas-based connectomes does not require construction of any intermediate high-resolution connectomes and is a fast operation relative to the time required to perform whole-brain tractography. Thus, while the benefits of spatial smoothing for atlas-based connectomes were modest, we recommend including CSS in future connectome mapping workflows.

### 4.6. CSS and principals of spatial smoothing

Our proposed connectome spatial smoothing approach is an extension of spatial signal smoothing to networks, and hence, fundamental concepts within the domain of spatial smoothing are applicable to CSS. For instance, from a signal processing perspective, the matched filter theorem states that spatial smoothing by an appropriate Gaussian kernel equalizes the voxel-wise standard deviation and, in turn, yields an optimal sensitivity to detect effects of unknown extent [77, 78]. Additionally, with regards to single-subject inference, such smoothing facilitates the application of multiple comparison correction using random field theory [77–79] and finally, smoothing mitigates residual anatomical variability of individuals at the group-level. These concepts are equally applicable to CSS, wherein, a matrix multiplication with a smoothing kernel achieves a similar purpose to convolution of image data with a 3D spatial smoothing kernel; as a result, CSS can be utilized to (i) maximize connectivity SNR through appropriate filter selection, (ii) improve single-subject inference, and (iii) improve the reliability of group level connectivity analyses. This poses an interesting future research direction to explore the benefits of CSS for whole-brain high-resolution network inference in which voxel-wise approaches [31, 79–81] are combined with network-based approaches [30].

### 4.7. Broad applicability of CSS

This study exclusively evaluated the merits of CSS for a measure of streamline count extracted from structural connectivity. However, the applicability of CSS extends beyond streamline counts to other diffusion-derived measures of connectivity, such as connection density [82], mean streamline length, and values sampled from quantitative diffusion model measures, e.g. the mean fractional anisotropy [83]. We note however that some care would be necessary if applying the method to metrics where the absence of a connection and a value of zero should not be treated equally.

Furthermore, its important to note that CSS is also applicable to brain networks derived from other imaging modalities. Functional connectivity (FC) generated from fMRI time-series data using e.g. Pearson’s correlation is one such example. Interestingly, CSS formulation can be applied to fMRI time-series to construct an equivalent smoothed FC with a moderate improvement in computational performance relative to traditional fMRI smoothing approaches (see Supplementary Information Section S.6. Smoothing functional connectivity with CSS for detail). This equivalence presents CSS as a fundamental generalization of image spatial smoothing that extends spatial smoothing to connectivity matrices.

### 4.8. Limitations

The main aim of this study was to evaluate the benefits of CSS for both high-resolution and atlas connectomes, but not to set a precedent for connectome construction decisions. Future work is needed to establish best practices in mapping high-resolution connectomes.

Following an earlier implementation [32], this study used a nearest endpoint assignment approach to map streamline endpoints to high-resolution vertices. The connectome construction process could be made more robust by e.g. terminating streamlines more precisely as they transition the grey-white interface surface and assigning them to the nearest vertex on that surface. However, the extent of the potential benefit and the computational complexity of implementing this procedure needs to be evaluated in future works.

Furthermore, it is important to note that the areal inequalities present in the fsLR-32k surface mesh can impose biases in connectome construction and alternative approaches could mitigate this limitation [33, 37] (see Supplementary Information Section S.7. Areal inequalities of the surface mesh for detail). While evaluating ramifications of such inequalities falls beyond the scope of this paper, future research should study the implications of these inequalities and propose appropriate correction strategies to alleviate areal connectivity biases.

### 4.9. Concluding remarks

In this study, we developed a novel formalism for spatial smoothing of structural connectivity matrices and demonstrated the wide-ranging benefits of connectome smoothing. Our results indicate that CSS with different kernel FWHMs and truncation thresholds significantly impacts various characteristics of structural connectivity matrices. In high-resolution connectomes, smoothing up to 3-6mm FWHM was deemed favorable, though the choice of smoothing parameters imposes a trade-off between reliability and individual identifiability. We provided recommendations for smoothing parameter choices that achieve a compromise between reliability and identifiability. Our connectome smoothing method and associated recommendations can be incorporated into future structural connectivity mapping pipelines, enabling more reliable and better powered connectome analyses. Moreover, high-resolution structural connectivity overcomes the known uncertainty and ambiguity in determination of brain parcellation, and so will be a powerful analysis framework moving forward; our demonstrated and evaluated smoothing framework is an essential tool in facilitating such, and we have made reasonable recommendations for how others can use it.

## Data and code availability

All imaging data used in this study was sourced from the Human Connectome Project (HCP) (www.humanconnectome.org). The bash scripts used to perform tractography using MRtrix3 [49] (www.mrtrix.org), as well as all Python code required to perform CSS and map smoothed connectomes at either the resolution of vertices or an atlas, are provided in our git repository. This code repository can be accessed from github.com/sina-mansour/connectome-based-smoothing. Additionally, to facilitate future research and promote open transparent practices in code-sharing [84–86], the codes for smoothing connectomes at high-resolution and atlas-resolution are released as a standalone python package [87].

## Acknowledgments

Data were provided by the Human Connectome Project, WU-Minn Consortium (Principal Investigators: David Van Essen and Kamil Ugurbil; 1U54MH091657) funded by the 16 NIH Institutes and Centers that support the NIH Blueprint for Neuroscience Research; and by the McDonnell Center for Systems Neuroscience at Washington University. The data analysis was supported by SPARTAN High Performance Computing System at the University of Melbourne [88], and also supported by use of the Melbourne Research Cloud (MRC) providing Infrastructure-as-a-Service (IaaS) cloud computing to the University of Melbourne researchers through the NeCTAR Research Cloud, a collaborative Australian research platform supported by the National Collaborative Research Infrastructure Strategy. S.M.L. is funded by a Melbourne Research Scholarship. R.S. is supported by fellowship funding from the National Imaging Facility (NIF), an Australian Government National Collaborative Research Infrastructure Strategy (NCRIS) capability. A.Z. was supported by a senior research fellowship from the NHMRC (APP1118153).

## Author contributions

**S.M.L**.: Conceptualization, Methodology, Formal analysis, Data curation, Software, Writing - original draft, Writing - review & editing **C.S**.: Conceptualization, Writing - original draft, Writing - review & editing **R.S**: Conceptualization, Writing - original draft, Writing - review & editing **A.Z**: Supervision, Conceptualization, Writing - original draft, Writing - review & editing

## Supplementary Information

### S.1. CSS for connectivity mapped on vertices vs. faces

High-resolution connectivity maps are defined as spatial networks on the cortical surface mesh. Previous research represents the nodes of this network using either the vertices [32, 34, 35, 39, 46, 47] or the faces (triangles) [33, 36, 37] of the surface mesh. While this study implemented CSS on a vertex-based high-resolution network, in this section, we demonstrate the duality between vertex-based and face-based connectivity representations and show that CSS could alternatively be applied on a face-based connectome representation. In the ensuing section, we first propose a primal-dual relationship that can be used to translate the vertex-based CSS definitions (primal representation) to a face-based CSS definition (dual representation).

#### S.1.1. Primal-dual representation

In the primal representation, the cortical surface is represented by a 3-dimensional triangular surface mesh defined by a set of vertices, and the triangulation scheme explaining how the surface curvature is constructed over vertices. This is the default representation normally provided in tools such as FreeSurfer. This space contains the following primal elements: i) a set of 0-dimensional coordinates of points called vertices, ii) a set of 1-dimensional line segments called edges which connect pairs of vertices together, iii) a set of 2-dimensional triangles called faces which consist of 3 edges each, and the collection of these triangles form a 3-dimensional surface representation.

In the dual representation, each of these primal elements are mapped to a complementary dual element which form a dual 3-dimensional structure (a 2-simplex mesh [89]) such that a unique (1-1) mapping between the primal and dual representations is possible:

- For every face in the primal, a vertex (0-dimensional element) is defined in the dual space positioned at the coordinate of the center of mass for that face. Along with this coordinate information, the normal vector of the primal face is also attributed to the dual vertex (this information is required to reconstruct the primal representation from the dual representation).
- For every edge in the primal, an edge (1-dimensional element) is defined in the dual space that connects the dual vertices associated to the primal triangles neighboring the primal edge. In other words, the dual edges connect the centers of mass of the neighboring primal faces.
- As a result, for every vertex in the primal, a dual representation is created by the set of dual representations of all primal edges connected to that primal vertex (a 2-face [90]). Basically, the dual representation is created by connecting the centers of mass of all triangles that the primal vertex is neighboring.

This forms a dual representation of the primal 3-dimensional surface in the form of a 2-simplex mesh (i.e. a 3-dimensional mesh in which every vertex is connected to 3 other vertices). This primal-dual representation is visualized on an example surface mesh in figure S1. A more detailed explanation of this primal-dual representation can be found elsewhere [89–91].

**Fig. S1.**
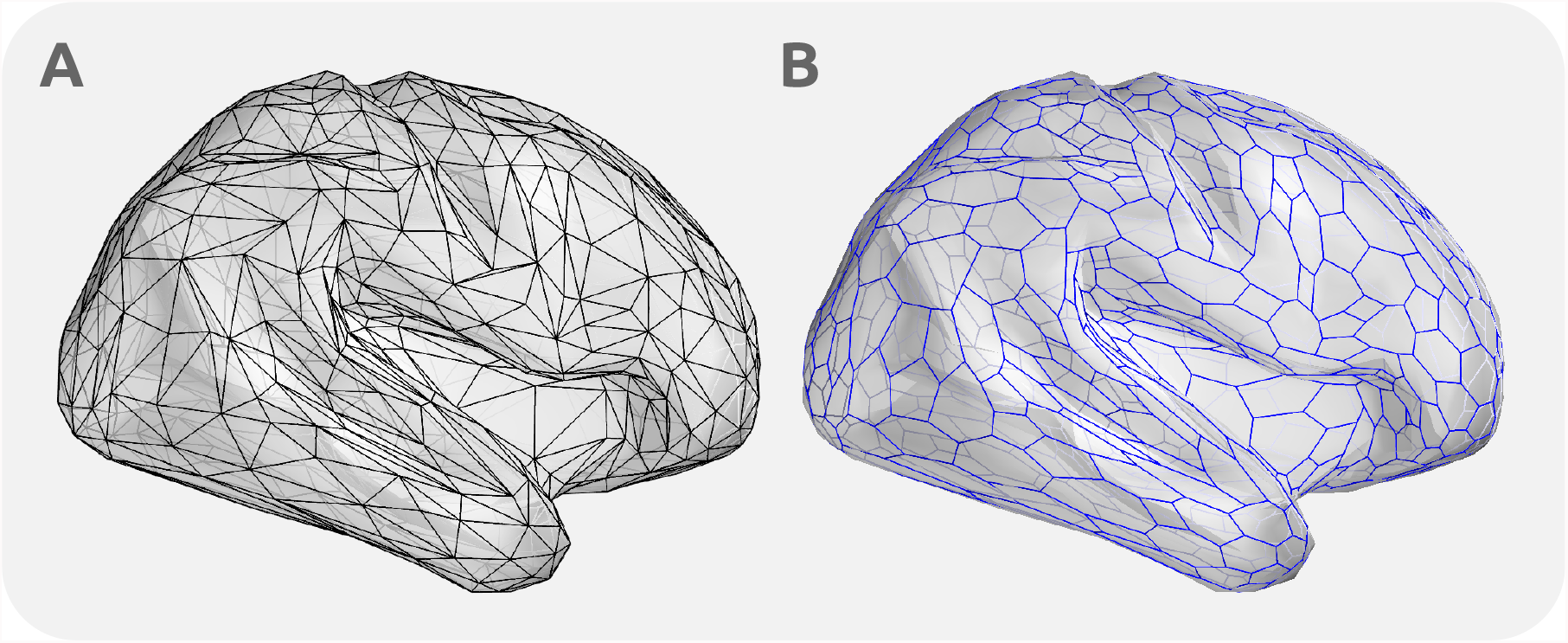
The primal-dual representation of a sample 3-dimensional surface mesh. A simplified surface mesh was created by quadric metric error decimation of the inflated right cortical surface. (A) The primal surface representation is visualized by black edges connecting primal vertices to form the primal faces. (B) The dual representation is visualized by blue edges connecting the dual vertices (located at the centers of mass of primal faces) and forming the dual polygons.

The dual representation provides a geometry to compute a spatial Gaussian smoothing kernel for face-based connectomes. This kernel is defined according to the geodesic distances between the dual vertices encoding primal faces in the dual representation. Hence, if high-resolution connectivity were to be mapped on a network of faces, this alternative smoothing kernel could be used to perform CSS on the face-based connectome.

In summary, high-resolution maps, when considered across all modalities and data formats used, are more commonly defined vertex-wise. However, for structural connectivity mapping via tractography, streamline intersections with faces arguably provide a more suitable native representation. Nevertheless, the natural duality between vertex- and face-based representations facilitates conversions of methods tailored for one to be applied to the other. In this study, we decided to prioritize the multimodal consistency of the mapped high-resolution connectomes by using a vertex-based mapping approach.

### S.2. Considerations for endpoint assignment

In this study we used a nearest-vertex check for endpoint assignment to high-resolution surface vertices (fsLR-32k). After tractography and removal of streamlines ending far (> 2mm) from cortical vertices, each streamline endpoint was assigned to its closest vertex according to 3D euclidean distance metric. Here, we elaborate on the implications of this choice and provide a comparison to other assignment approaches. An implication of this selection for endpoint assignment method is the potential ambiguity in assignment boundaries. For the sake of comparison, high-resolution structural connectome studies that assign endpoints to faces simply check for the intersection of streamline at each endpoint with a surface mesh face. This provides a clear areal boundary definition of assignment borders (Figure S2.A).

**Fig. S2.**
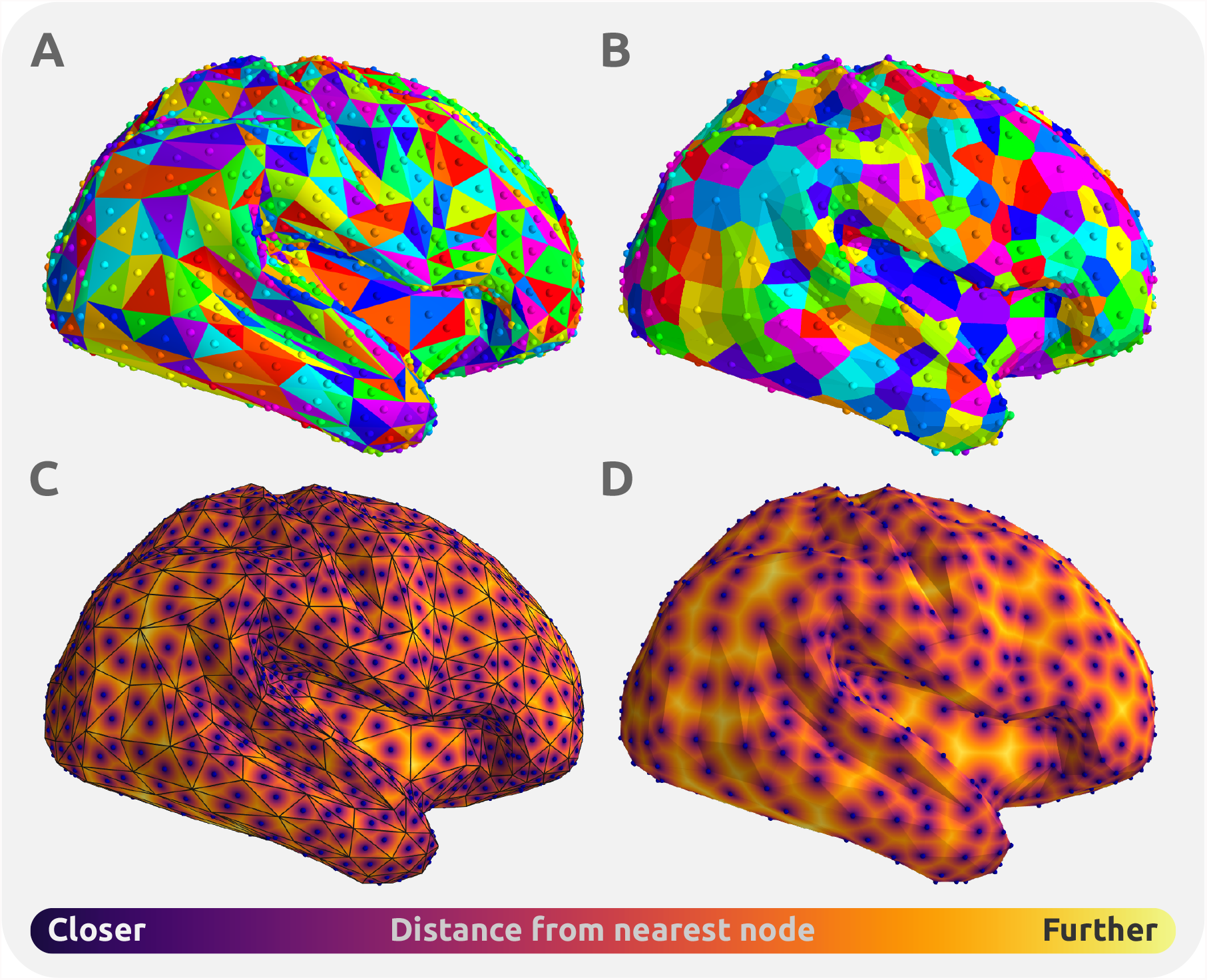
Endpoint assignment areal boundaries. A simplified surface mesh created by quadric metric error decimation of the inflated right cortical surface was used as an example. (A) The face-based assignments are shown on the colored surface. The center of mass for each face is colored independently; the same color is also used to delineate the surface boundaries for streamline assignment with the intersection based approach. This approach assigns each streamline to the face it intersects with. (B) The vertex-based endpoint assignment used in this study is used to generate a similar coloring. Every vertex is independently colored, a Voronoi tessellation was projected on the surface mesh to delineate assignment boundaries for every vertex. (C) & (D) The distance of points on the surface mesh to their closest node is visualized such that higher distances are assigned a warmer color. The distances for the face intersection approach do not necessarily follow the assignment boundaries delineated by faces; Whereas, the nearest vertex assignments are designed to follow the distance boundaries by definition.

In the following, we discuss how the consequences of our endpoint assignment approach can be translated to a similar areal boundary definition resolving assignment ambiguities.

Fundamentally, since the endpoints are treated as coordinates assigned to the closest vertices on the surface mesh, a Voronoi tessellation can be computed to evaluate the inclusion boundaries to assign an arbitrary endpoint to a vertex. This Voronoi tessellation can be projected to the surface mesh to delineate the assignment boundaries on the surface mesh. Figure S2.B presents how this area boundary could be defined for vertex-based streamline assignments on the triangular surface meshes. Furthermore, Figures S2.C,D projects the Euclidean distance from the closest node (vertex/face) along the surface manifold. These results suggest that the endpoint assignment by intersection approach previously used in face-based connectomes will not necessarily assign a streamline to the closest node (according to the distance metric). Hence, the two endpoint assignment approaches are different as one checks for the intersection of a streamline with a face on the surface, and the other assesses proximity to the closest vertex. Nonetheless, both approaches define a clear areal boundary along the surface mesh which can be used to evaluate streamline inclusion. These alternative approaches could also be theoretically expanded to the dual representation of the simplex mesh. In other words, if the aim is to generate vertex-based connectivity with an intersection based endpoint assignment, or to generate face-based connectivity with a nearest face assignment constraint, the dual surface representation can be used.

### S.3. Thresholding radius

This section provides the mathematical rationale behind the relationship between *R, σ* (or alternatively FWHM), and *ε* presented in Equation 6. The truncation radius *R* was formulated as a function of *σ* and *ε* such that the proportion of signal loss for a 2-dimensional Gaussian kernel with the strength of *σ* truncated at a radius of *R* is equal to *ε*. Given that the Gaussian kernel was defined such that its total cumulative density is unity 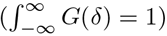, the relationship between the smoothing parameters can be defined by the following integration over the 2-dimensional surface area:

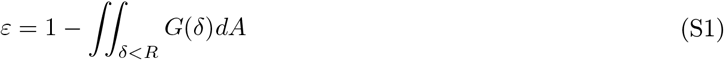

This integration can be solved in polar coordinates by the following closed form equation:

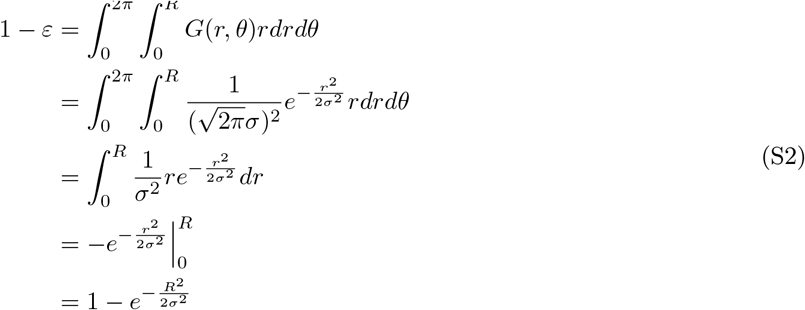

And this can be used to describe *R* as a function of *σ* and *ϵ*:

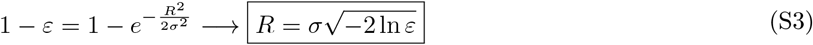

And given the relationship between FWHM and 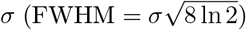, this equation can be rewritten based on FWHM:

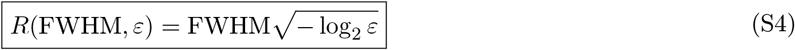

### S.4. CSS for atlas-based connectivity

In the main text, it was briefly mentioned that mapping the high-resolution connectivity is not necessary for smoothing the connectivity matrices at an atlas resolution: alternatively, a smoothed version of an atlas-based connectivity matrix can be derived from a soft parcellation, which is derived by applying spatial smoothing to the parcels of the brain atlas (and normalizing each vertex to a unity sum of parcel memberships). In this section, we provide the formal proof of this equivalence: first, downsampling a high-resolution connectivity matrix to an atlas-based connectome matrix is formulated by linear algebraic formulations; these formulations are then used to complete a formal proof of the equivalence.

Following the prior nomenclature, *A* is a *v* × *v* matrix denoting the high-resolution connectivity matrix where *v* is the number of vertices. According to Equation 4, the smoothed high-resolution connectivity matrix *A*_*s*_ is calculated as follows:

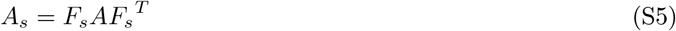

Where *F*_*s*_ is a *v* × *v* column-normalized spatial smoothing kernel. A formal notion of a brain atlas can be denoted by *p* × *v* matrix P, where *p* is the number of brain regions in the atlas. Elements *P* (*i, j*) encode the relationship between vertex/voxel *v*_*i*_ and region *p*_*j*_.

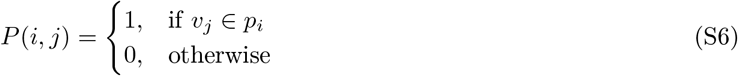

An atlas-resolution connectome *A*_*p*_ is a *p* × *p* matrix, which is normally mapped from an atlas parcellation such that elements *A*_*p*_(*i, j*) encode the aggregate contribution from those streamlines for which one endpoint is assigned to region *p*_*i*_ and the other endpoint is assigned to region *p*_*j*_ (showing here the streamline count for simplicity):

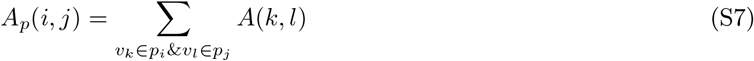

This notion can be formalized by the following matrix representation which can be used to derive *A*_*p*_ from *A* and *P* :

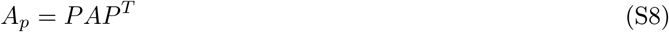

Hence, the element *A*_*p*_(*i, j*) counts the overall connectivity between regions *p*_*i*_ and *p*_*j*_ by adding all high-resolution connectivity edges between them. Equations S5 and S8 yield the following definition for the smoothed atlas-based connectivity *A*_*sp*_:

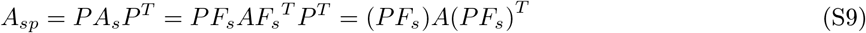

The matrix *PF*_*s*_ can thus be treated as a *p* × *v* weighted soft parcellation map, i.e. a non-binary brain atlas. This soft parcellation can be used to generate smoothed connectomes based on an atlas parcellation (each streamline contributes to many connectome edges, based on all parcels with non-zero densities at both end-points) (see Equation S8). A key benefit of this approach is that it obviates the need to create computationally cumbersome high-resolution connectomes as an intermediate step in construction of lower-resolution connectome matrices. A different approach to compute this soft parcellation, that additionally does not necessitate computation of high-resolution smoothing matrix *F*_*s*_, is further described in the ensuing sections.

#### S.4.1. Column normalization

To describe the soft parcellation *PF*_*s*_, a formal definition of normalizing every column should first be defined. Column normalization of an *l × m* matrix *B* can be defined by the matrix multiplication of *B* with a diagonal norm matrix constructed from column sums.

##### Definition S.1.

⟨|*B*|⟩ *denotes an m* × *m diagonal column norm matrix constructed from B where* ⟨|*B*|⟩ (*i, i*) *is the sum of the elements of the ith column in B:*

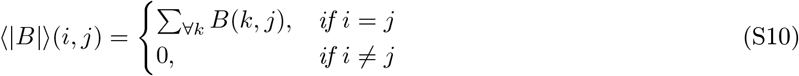

*Hence*,

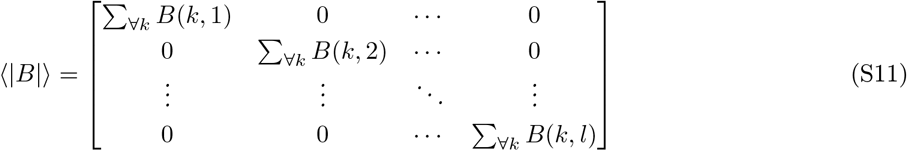

And a consequence of Definition S.1 is the statement in the next corollary.

##### Corollary S.1.1.

*Let z*_*i*_ ∈ ℝ^*i*^ *denote the vector of ones, i*.*e. all i vector elements equal 1. The following is true for any diagonal norm matrix:*

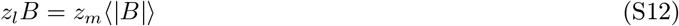

Both sides of the equation above compute the column sums of *B*. Column normalization can be formally defined by the following theorem.

##### Theorem S.1.

*The row normalization is a matrix transformation of an l* × *m matrix B to an l*× *m normalized matrix N* (*B*), *such that the sum of every column in N* (*B*) *is equal to 1, i*.*e. z*_*l*_*N* (*B*) = *z*_*m*_. *N* (*B*) *can be derived by the following matrix multiplication:*

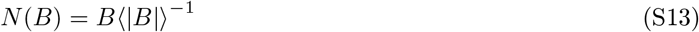

*Proof*. Corollary S.1.1 can be used to prove *z*_*l*_*N* (*B*) = *z*_*m*_:

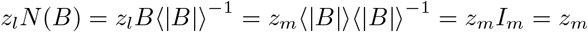

□

Where *I*_*m*_ is the *m*× *m* identity matrix. The following remarks are a consequence of the aforementioned definitions and theorems.

##### Remark.

*The diagonal norm matrix of a brain atlas parcellation* ⟨|*P*|⟩ *is the v* ×*v identity matrix I*_*v*_, *as every vertex belongs to a single atlas region and thus the sum of any column of P equals 1:*

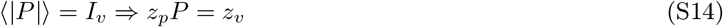

*Thus, for any arbitrary v* × *v matrix X:*

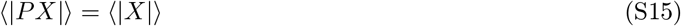

⟨|*PX*|⟩, *by definition, is a diagonal matrix:*

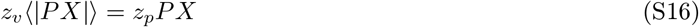

*and from Equation S14 we know that:*

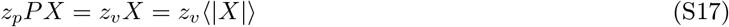

*Therefore*, ⟨|*X*|⟩ *is the same diagonal matrix as* ⟨|*PX*|⟩. *In other words, the sum of the columns of PX is equal to the sum of the columns of X*.

##### Remark.

*The normalized high-resolution smoothing kernel F*_*s*_ *is defined from column normalization of the Gaussian kernel smoothing weights matrix F*_*G*_ *(from Equation S13):*

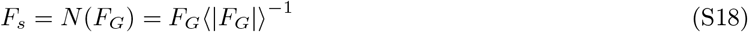

*Where F*_*G*_ *is a symmetric v* ×*v matrix yielded from the truncated Gaussian function calculated upon the surface mesh:*

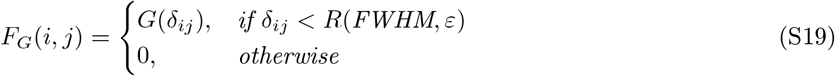

#### S.4.2. Smoothed brain atlas

Equation S9 showed that a smoothed soft parcellation *P*_*s*_ = *PF*_*s*_ can be used to directly derive smoothed atlas connectivity maps from tractography. In this section, a formal proof will be provided for the following statement:

##### Theorem S.2.

*The smoothed soft parcellation P*_*s*_ = *PF*_*s*_ *can be computed in the absence of F*_*s*_, *by separately smoothing every row of P, followed by normalizing every column of the smoothed parcellation:*

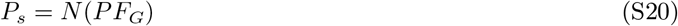

*Proof*. Using the previously derived equations, we prove that *P*_*s*_ = *N* (*PF*_*G*_):

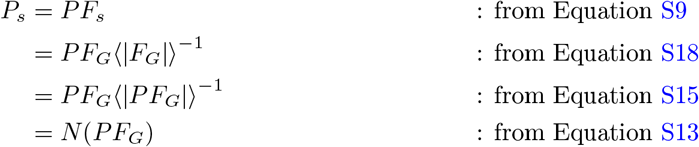

□

The proof above confirms that structural connectivity based on a parcellation atlas, incorporating CSS, can be constructed directly from a tractogram and soft parcellation, without necessitating computation of either the high-resolution smoothing matrix or the high-resolution connectome. To smooth an atlas-resolution connectome, the brain atlas *P* should first be transformed to a normalized smoothed soft parcellation *P*_*s*_ = *N* (*PF*_*G*_). *PF*_*G*_ is equivalent to independently smoothing the binary representation of each parcel, while the normalization of such ensures that the sum of parcel memberships of every vertex is 1. Hence, the soft-parcellation *P*_*s*_ can be computed by spatial smoothing and then be directly combined with the tractogram to produce a connectome: each streamline endpoint may have non-zero attribution to multiple parcels, and the contribution of the streamline to the connectome is therefore distributed across the set of edges associated with those two sets of parcels. This constitutes an approach to apply CSS on atlas-resolution connectomes that does not require any high-resolution connectomic computations.

### S.5. Replication of ROC curve estimates

The computation of ROC curves reported in the manuscript relied on the assumption of a fixed false omission rate 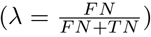. To ensure that the findings were not biased by the selected value for *λ*, the same analyses was repeated for a range of plausible values of *λ* ∈ 10%, 1%, 0.1%}. Fig. S3 presents the results of this evaluation. The findings indicate that CSS increases the sensitivity of the statistical analyses and the inference power, particularly for connectomes mapped from probabilistic tractography, regardless of the selection made for the false omission rate *λ*.

**Fig. S3.**
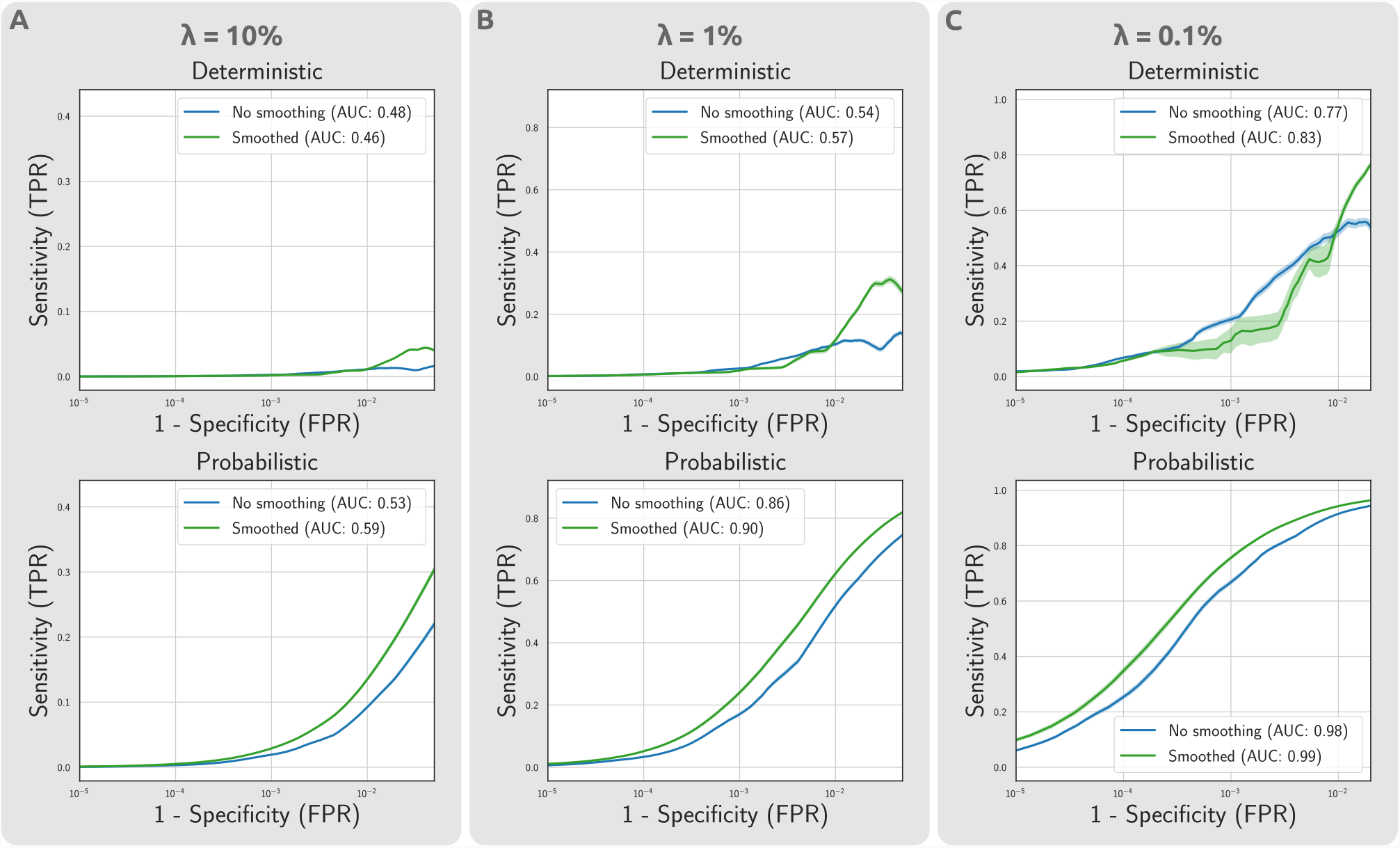
Impact of CSS on statistical power of mass univariate testing on atlas-based connectomes, for different false omission rate assumptions. The estimated ROC curves demonstrate the respective changes in sensitivity and specificity of the suprathreshold edges at different effect thresholds. The analyses was repeated across a range of false omission rates to ensure the robustness of findings with regards to parameter selection. While absolute model performance estimate quantified by AUC depends on the false omission rate assumption, the findings with regards to the relative performance of methods are consistent across different false omission rates.

### S.6. Smoothing functional connectivity with CSS

In this section, we evaluate how the mathematical formulations of CSS could alternatively be used for smoothing functional connectivity (FC) matrices mapped at the resolution of a brain atlas. Formally, an fMRI time-series can be denoted by a *v* ×*t* matrix *C*, where *t* is the number of time-points in the fMRI data. Essentially, every row of matrix *C* contains the time-series of a single node in high-resolution.

Traditionally, to map smoothed FC matrices, this data is first smoothed with a Gaussian spatial smoothing kernel. Borrowing from CSS formulations, the smoothed time-series data *C*_*s*_ can be computed by a matrix multiplication of the time-series with the connectome smoothing kernel:

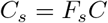

Next, the spatially smoothed time-series is downsampled to the resolution of a brain atlas:

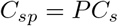

Finally, computing the Pearson’s correlation between all pairs of regions (rows of *C*_*sp*_) generates the final FC matrix:

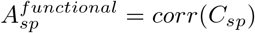

Where, *corr* denotes the function that computes row-wise correlations and 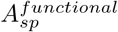 denotes the final FC matrix. Hence, the traditional procedure for mapping smoothed FC matrices can be represented by the following CSS notation:

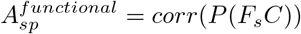

The atlas based simplification of CSS (see Supplementary Information Section S.4. CSS for atlas-based connectivity) could thus be used to speed up the traditional FC mapping pipelines. Instead of smoothing the fMRI time-series followed by downsampling to the atlas resolution, the unsmoothed fMRI time-series could be directly downsampled with a smoothed soft parcellation (*P*_*s*_ = *PF*_*s*_):

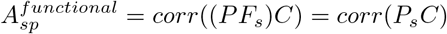

This alternative approach inspired by atlas-based CSS provides an improvement in computational complexity of smoothed FC mapping (from *O*(*pv*^2^ + *v*^2^*t*) to *O*(*pv*^2^ + *pvt*) for a single FC). Furthermore, if smoothed FC is mapped for multiple individuals, *P*_*s*_ can be computed once for the group average template space. Hence, the amortized time complexity is reduced from the traditional complexity of *O*(*pv*^2^ + *v*^2^*t*) to 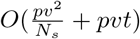, where *N*_*s*_ is the number of individuals for whom FC is being mapped. Although this relatively moderate improvement may not justify modification of traditional FC mapping pipelines, it demonstrates how CSS can be viewed as a fundamental generalization of traditional spatial smoothing approaches to a connectivity domain.

### S.7. Areal inequalities of the surface mesh

This study used the vertices from fsLR-32k atlas to delineate high-resolution structural connectivity nodes. Using this delineation is beneficial due to its alignment to other properties extracted from neuroimaging scans, such as cortical thickness, curvature, functional activation and connectivity, and cortical atlases. More importantly, these surfaces are all registered to the same standard sphere and hence provide a 1-1 mapping of vertices across individuals. Nevertheless, it is important to highlight the shortcomings of this surface mesh in the context of high-resolution connectomic studies.

As this surface was generated to closely follow the individual’s curvature of the cortical WM surface for volumetric preservation, the vertices are not necessarily distributed equidistantly on the cortical surface [37] (similarly, as per Section S.1.1, the faces are not equiareally distributed on the surface). In this section we evaluate the distance and areal inequalities of the fsLR-32k mesh to highlight the connectivity biases created by such inequalities. High-resolution connectivity studies may benefit from correction of such biases to ensure that any results reported are not impacted/contaminated by these inequalities.

In order to investigate the effect of these biases, we computed two measures to indicate distance and area inequalities observed in the fsLR-32k mesh. First, we computed the average distance between every vertex and all of its direct neighbors, i.e. vertices sharing an explicit primal surface edge. Additionally, we computed a vertex-based area measure by distributing the area of faces across the surrounding vertices on the surface. This lumped vertex area quantified the areal inequalities of the surface mesh that could impact vertex-based connectivity. Fig. S4.A,B shows the distribution of these measures for the surface of an exemplar individual. Additionally Fig. S4.C,D shows the same measures averaged across all individuals (N=42). These results suggest the existence of distance and areal inequalities along the fsLR-32k mesh that are mostly correlated such that regions with higher intervertex distances contain triangles with higher areas. The group average results indicate the inherent distance and areal biases of the fsLR-32K mesh, such that regions further from the center of the original sphere (i.e. frontal, occipital, and temporal regions) tend to stretch more and thus have a higher area and intervertex distance compared to regions closer to the center of original sphere (i.e. somatosensory and motor cortices as well as medial and insular regions).

Such inequalities may introduce inhomogeneity in the mapped connectomes and impact high-resolution connectivity studies. To demonstrate the potential impacts of these inequalities on connectivity measures, Fig. S4.E,F shows the correlation between weighted degree distribution (i.e. nodal strength) in the unsmoothed connectome and the intervertex distance and vertex area variations. These results indicate a significant yet modest association between both measures of distance (*r* = 0.089, *p* = 0.025) and area (*r* = 0.124, *p* = 0.007) with nodal strength (significance was evaluated by a non-parametric spin test [92]). This indicates that regions with higher intervertex distance/area tend to have more streamlines assigned to the nodes within them. An earlier high-resolution study [33] have suggested an areal normalization approach by which the high-resolution connectivity edges indicating streamline counts were normalized by areas of the node pairs. The normalization replaces every edge weight *e*_*ij*_ indicating streamline count between nodes *v*_*i*_ and *v*_*j*_ with a normalized edge weight 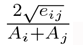, where *A*_*i*_ and *A*_*j*_ denote the areas associated with nodes *v*_*i*_ and *v*_*j*_. We tested to see if this normalization method could reduce/remove the distance and areal biases in nodal strength (Fig. S4.G,H). Our results indicate that after performing the suggested normalization, nodal strength was still significantly associated with both distance (*r* = − 0.113, *p* < 0.001) and area (*r* = 0.105, *p* < 0.001). The only difference was in the direction of the association, such that vertices with higher distance/area tended to have a lower normalized connectivity. Future studies are needed to investigate areal and distance biases and propose alternative normalization approaches that could remove/reduce the impacts of these biases.

Another alternative solution used in previous studies is using isotropic remeshing to reduce the extent of areal inequalities in the native surface mesh [37]. Nevertheless, implementing isotropic remeshing would require an extra resampling step in order to carry vertex-wise comparisons across subjects. The extent of benefits of this alternative approach was not evaluated in this study and requires future investigation.

**Fig. S4.**
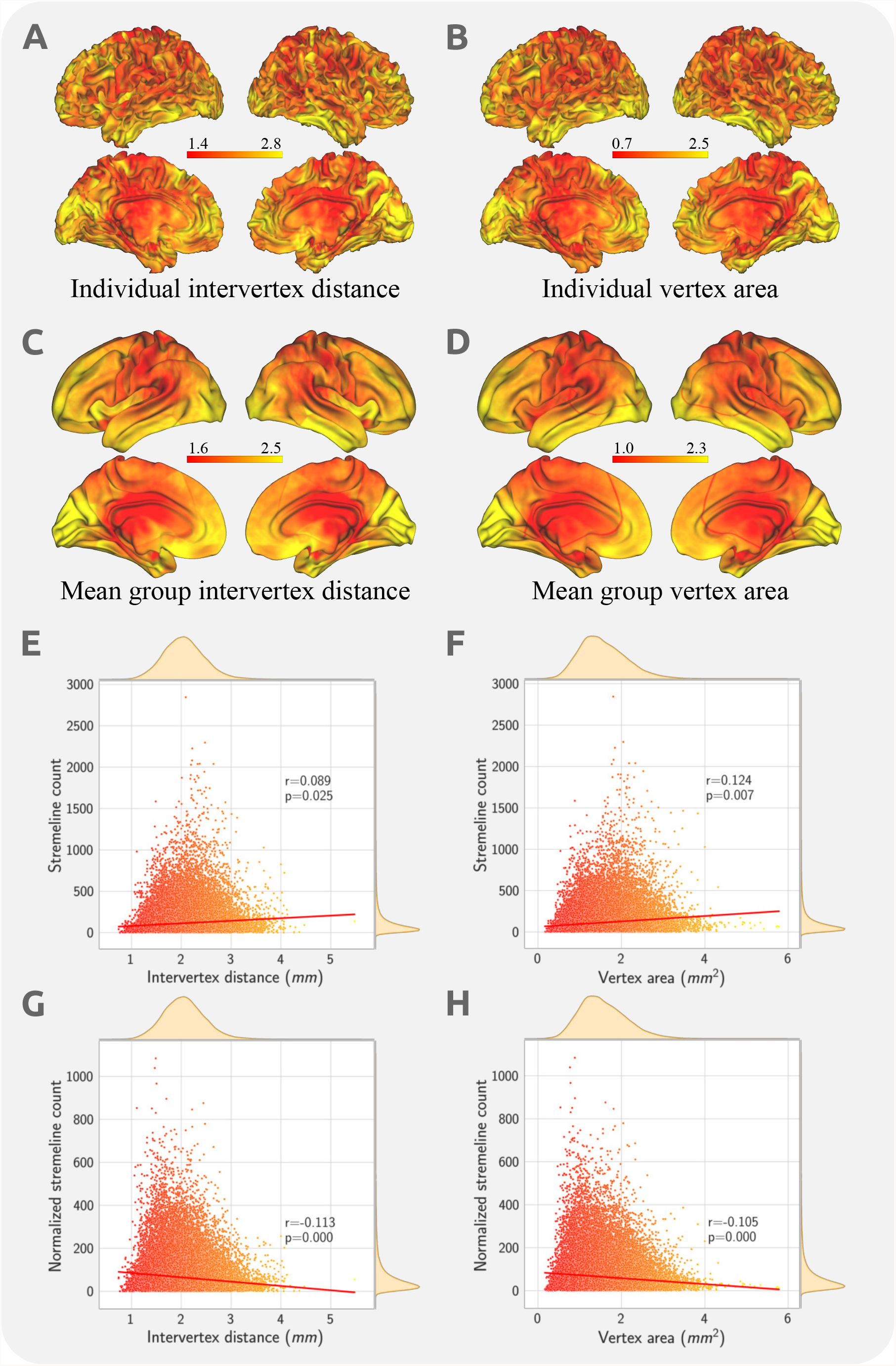
Impact of distance and areal inequalities on high-resolution connectivity mapping. Panels (A) and (B) respectively show the intervertex distance and lumped vertex area inequalities projected over the same individual’s WM fsLR-32K surface. Panels (C) and (D) show the inequalities averaged across the group (N=42) projected on the group-average WM surface. The group average maps indicate group-level general biases that the frontal, occipital, and temporal regions are more stretched. Hence, vertices in these regions generally have higher associated area and intervertex distance. Panels (E) and (F) provide scatter plots with marginal distributions indicating the associations between distance and areal inequalities (shown in panel A,B) and nodal streamline count. The results indicate a modest yet significant impact of areal and distance inequality on streamline count. Statistical significance was evaluated by a non-parametric spin test. Panels (G) and (H) provide similar scatter plots for normalized streamline counts and show that the biases remain even after normalization.

